# Genomic analysis of carbapenemase-encoding plasmids from *Klebsiella pneumoniae* across Europe highlights three major patterns of dissemination

**DOI:** 10.1101/2019.12.19.873935

**Authors:** Sophia David, Victoria Cohen, Sandra Reuter, Anna E. Sheppard, Tommaso Giani, Julian Parkhill, the European Survey of Carbapenemase-Producing Enterobacteriaceae (EuSCAPE) Working Group, the ESCMID Study Group for Epidemiological Markers (ESGEM), Gian Maria Rossolini, Edward J. Feil, Hajo Grundmann, David M. Aanensen

## Abstract

The incidence of *Klebsiella pneumoniae* infections that are resistant to carbapenems, a last-line class of antibiotics, has been rapidly increasing. The primary mechanism of carbapenem resistance is production of carbapenemase enzymes, which are most frequently encoded on plasmids by *bla*_OXA-48-like_, *bla*_VIM_, *bla*_NDM_ and *bla*_KPC_ genes. Using short-read sequence data, we previously analysed genomes of 1717 isolates from the *K. pneumoniae* species complex submitted during the European survey of carbapenemase-producing *Enterobacteriaceae* (EuSCAPE). Here, we investigated the diversity, prevalence and transmission dynamics of carbapenemase-encoding plasmids using long-read sequencing of representative isolates (*n*=79) from this collection in combination with short-read data from all isolates. We highlight three major patterns by which carbapenemase genes have disseminated via plasmids. First, *bla*_OXA-48-like_ genes have spread across diverse lineages primarily via a highly conserved, epidemic pOXA-48-like plasmid. Second, *bla*_VIM_ and *bla*_NDM_ genes have spread via transient associations of diverse plasmids with numerous lineages. Third, *bla*_KPC_ genes have transmitted predominantly by stable association with one clonal lineage (ST258/512) despite frequent mobilisation between pre-existing yet diverse plasmids within the lineage. Despite contrasts in these three modes of carbapenemase gene spread, which can be summarised as using one plasmid/multiple lineages, multiple plasmids/multiple lineages, and multiple plasmids/one lineage, all are underpinned by significant propagation along high-risk clonal lineages.

## Introduction

The incidence of infections due to carbapenem-resistant *Enterobacterales* (CRE) is rapidly rising, posing a major challenge to public health globally (WHO, 2017). Indeed, carbapenem-resistant *Klebsiella pneumoniae*, the most clinically significant member of CRE, was recently highlighted as the fastest-growing resistance threat in Europe in terms of number of infections and attributable deaths (Cassini et al. 2019).

The largest subset of CRE, the carbapenemase-producing *Enterobacterales* (CPE), hydrolyse carbapenems and other beta-lactam antibiotics using diverse types of beta-lactamase enzymes called carbapenemases (David et al. 2019). Genes encoding these carbapenemases are usually located on plasmids which can transmit vertically along clonal lineages as well as horizontally between different strains and species (Mathers et al. 2011; Martin et al. 2017). Within plasmids, carbapenemase genes are also frequently associated with smaller mobile genetic elements such as transposons and mobile gene cassettes inserted into integrons, extending their recombinatory capability to multiple nested levels (Sheppard et al. 2016).

Next-generation sequencing using short-read technologies has vastly improved our ability to unravel the complexities of infectious disease epidemiology. In particular, it has enabled genomic surveillance of high-risk bacterial lineages including tracking of their geographical dissemination (Aanensen et al. 2016; Domman et al. 2017; Harris et al. 2018; David et al. 2019). These surveillance approaches typically use differences in a defined chromosomal region (the “core genome”) that are determined by mapping sequence reads to a reference. However, advances in short-read sequencing have not enabled the same high-resolution tracking of plasmids since, typically being mosaic and recombinant, these usually require *de novo* assembly for accurate comparison. Unfortunately plasmid assemblies derived from short-read data are usually highly fragmented as a result of numerous repetitive elements (e.g. insertion sequences), and often cannot be distinguished from chromosomal sequences. Recently, these problems have been overcome by the advent of long-read sequencing, which now readily enables complete (or near-complete) and accurate resolution of plasmid sequences, particularly when the data are assembled together with short reads (Wick et al. 2017; George et al. 2017). This advance, coupled with the decreasing costs of long-read sequencing, renders large-scale plasmid comparisons increasingly feasible and brings the benefits of the sequencing revolution to bear also on the molecular epidemiology of plasmids.

Despite the rapidly growing databases of carbapenemase-encoding plasmid sequences, no study has systematically analysed the diversity of these plasmids in clinical isolates across a large, unbiased and geographically diverse sample collection. Previously, we analysed genomes of 1717 clinical isolates belonging to the *K. pneumoniae* species complex sampled from 244 hospitals in 32 countries during the European survey of CPE (EuSCAPE) (Grundmann et al. 2017; David et al. 2019). Six hundred and seventy-eight (39.5%) carried one or more of the *bla*_OXA-48-like_, *bla*_VIM_, *bla*_NDM_ and *bla*_KPC_ carbapenemase genes. All carbapenemase-positive isolates belonged to the *K. pneumoniae sensu stricto* species with the exception of one isolate each from *Klebsiella quasipneumoniae* and *Klebsiella variicola*. Here we investigated the diversity of carbapenemase-encoding plasmids amongst these isolates using combined long- and short-read sequencing of selected representatives. Furthermore, we explored the potential and limitations of using short-read sequence data obtained from all isolates, together with reference plasmids obtained from representatives, to assess the prevalence, distribution and transmission dynamics of carbapenemase-encoding plasmids across the wider European population. These analyses revealed three major patterns of plasmid transmission that have enabled widespread dissemination of carbapenemase genes.

## Results

### Diversity of the genetic contexts of carbapenemase genes among *K. pneumoniae*

Of 1717 *K. pneumoniae* species complex isolates submitted during the European Survey of CPE (EuSCAPE), we previously found that 249, 56, 79 and 312 carried *bla*_OXA-48-like_, *bla*_VIM_, *bla*_NDM_ and *bla*_KPC_ genes, respectively (David et al. 2019). Eighteen of these carried two genes. In this study, we first analysed the genetic contexts of these genes in short-read genome assemblies, considering this feature as a proxy for putative plasmid diversity. Assembly contigs containing each of the four carbapenemase genes were clustered into genetic context (GC) groups, based on the order and nucleotide similarity of genes flanking the carbapenemase gene (**Supplementary Tables 1** and **2**). Full contig sequences were used for this analysis. Contigs with fewer than four genes were excluded.

By this criterion, we identified 3, 10, 15 and 45 GC groups of isolates with different genetic contexts of *bla*_OXA-48-like_, *bla*_VIM_, *bla*_NDM_ and *bla*_KPC_ genes, respectively (**Table 1**). Overall, 184/696 (26.4%) of carbapenemase-carrying contigs could be unambiguously assigned to one of these groups. Assignment rates were higher for isolates carrying *bla*_VIM_ and *bla*_NDM_, and lower for those carrying *bla*_KPC_ and *bla*_OXA-48-like_. In particular, only 4/249 (1.6%) of *bla*_OXA-48-like_-carrying isolates could be assigned to a GC group due to the small size of the contigs, which typically carried only *bla*_OXA-48-like_ +/− *lysR* genes.

**Table 1.** Number of isolates assigned to different genetic context (GC) groups of the carbapenemase genes using short-read sequencing data. IQR – inter-quartile range.

| Carbapenemase gene | No. of isolates | Median no. of genes per short-read contig (IQR) | No. (%) of isolates with assigned GC group | No. (%) of isolates discarded from clustering* | No. (%) of isolates with ambiguous context** | No. of GC groups |
| --- | --- | --- | --- | --- | --- | --- |
| *bla*<sub>OXA-48-like</sub> | 249 | 2 (2-2) | 4 (1.6%) | 221 (88.8%) | 24 (9.6%) | 3 |
| *bla*<sub>VIM</sub> | 56 | 6 (2-7) | 39 (69.6%) | 16 (28.6%) | 1 (1.8%) | 10 |
| *bla*<sub>NDM</sub> | 79 | 15 (4-22) | 59 (74.7%) | 6 (7.6%) | 14 (17.7%) | 15 |
| *bla*<sub>KPC</sub> | 312 | 23 (18-35) | 82 (26.3%) | 0 (0%) | 230 (73.7%) | 45 |

\*Isolates were discarded from the clustering either because there were fewer than four genes on the carbapenemase-carrying contig, or if the carbapenemase gene was found in the assembly without a start or stop codon.
\*\*Isolates were designated an “ambiguous” context if the carbapenemase-carrying contig matched multiple, different (often larger) contigs.

We selected one isolate from each GC group for long-read sequencing, with the exception of one *bla*_OXA-48-like_ group and two *bla*_KPC_ groups for which representative isolates were unavailable. Furthermore, since the above-described process resulted in selection of only two *bla*_OXA-48-like_-carrying isolates, we also selected an additional eight. These had *bla*_OXA-48-like_-carrying contigs with ≥4 genes and, despite not clustering unambiguously into a single GC group, matched different combinations of other *bla*_OXA-48-like_-carrying contigs (**Supplementary Table 2**). They were therefore deemed the most likely to represent different plasmids amongst the remaining isolates. We also long-read sequenced another four *bla*_OXA-48-like_-carrying isolates which were positive for two carbapenemase genes and had been selected as representatives of *bla*_KPC_, *bla*_VIM_ or *bla*_NDM_ GC groups. Furthermore, one isolate selected as a representative of a *bla*_KPC_ GC group also harboured *bla*_VIM_. Finally, we long-read sequenced two *bla*_KPC_-carrying isolates from the same GC group to investigate possible within-hospital plasmid transfer, since they were submitted from the same hospital but belonged to different sequence types (ST).

### Long-read sequencing of representative isolates revealed that most carbapenemase genes were plasmid-borne

We assembled long-read sequencing data together with the previously obtained short reads for 79 isolates, encoding a total of 84 carbapenemase genes (**Supplementary Table 3**). The total number of contigs in the resulting hybrid assemblies ranged from 2-44 (median, 9). In 61/79 (77.2%) hybrid assemblies, the largest contig was ≥5Mb, indicating that all, or most, of the chromosomal sequence assembled into a single contig. The assemblies contained 1-8 plasmid replicons (median, 4), which are sequences used for defining plasmid incompatibility (Inc) groups (Carattoli et al. 2014). Multiple plasmid replicons were commonly found on the same contig, representing fusions between different plasmid types.

We found one copy of each carbapenemase gene in the hybrid assemblies. Five (3x *bla*_OXA-48-like_, 1x *bla*_KPC_, 1x *bla*_VIM_) were located on contigs ranging in size from 3.3-5.4Mb, each representing either a partial or putatively complete chromosomal sequence. The remaining 79 genes were located on contigs ranging in size from 2.5-313.6kb, which are hereafter described as putative plasmid sequences. Indeed, a plasmid origin is supported by the circularisation of 44 (55.7%) of these sequences, as well as the identification of plasmid replicons in 65 (82.3%). Of 11/79 (13.9%) putative plasmid sequences that could neither be circularised nor contained plasmid replicons, we found additional evidence of a plasmid origin for ten (see **Supplementary Note**).

### Dissemination of *bla*_OXA-48-like_ genes by rapid spread of pOXA-48-like plasmids across diverse lineages

Amongst the 14 *bla*_OXA-48-like_-carrying hybrid assemblies obtained, we found the carbapenemase gene in three chromosomal sequences (3.3-5.4Mb) and 11 putative plasmid sequences (2.5-149.6kb) (**Supplementary Table 3**). The two isolates sequenced as GC group representatives harboured *bla*_OXA-48-like_ on IncX3 and IncA/C2 plasmids, although we also found IncL/M(pOXA48) (*n*=3), IncL/M(pMU407) (*n*=1) and ColKP3 (*n*=3) plasmids carrying *bla*_OXA-48-like_ amongst the additional hybrid assemblies. Notably, three IncL/M(pOXA48) plasmids of 61.1-63.5kb showed high structural and nucleotide similarity to a well-described, 61.8kb plasmid, pOXA-48a from strain 11978 (Poirel et al. 2012), which belongs to the pOXA-48-like family (**Supplementary Figure 1;** see **Supplementary Note**).

We determined the prevalence of the different *bla*_OXA-48-like_-carrying plasmid sequences across all *bla*_OXA-48-like_-carrying isolates in the sample collection (*n*=249) by mapping the short sequence reads to each of the putative plasmid sequences obtained from the hybrid assemblies (see *Methods*). Importantly, this approach cannot reveal whether there have been insertions or rearrangements relative to the reference plasmid, or whether a particular resistance gene (in this case, *bla*_OXA-48-like_) is integrated into the same plasmid or located elsewhere, but nevertheless provides an indication of how much of each plasmid backbone is present.

Using this approach, we found that the IncX3, IncA/C2, IncL/M(pMU407) and ColKP3 plasmid sequences were found in full only rarely amongst all 249 isolates (**Figure 1A;** see **Supplementary Note**). In contrast, 204/249 (81.9%) isolates had short reads that mapped to ≥99% of the circularised 63.5kb IncL/M(pOXA48) plasmid from EuSCAPE_MT005 (and 221/249 (88.8%) to ≥90%). These comprised 202 isolates with the *bla*_OXA-48_ variant and two with *bla*_OXA-162_. For non-*bla*_OXA-48-like_-carrying isolates from the sample collection (*n*=1468), the median length of mapping to this plasmid was just 2.9% (interquartile range, 2.4-4.9%) while only 20/1468 (1.4%) mapped to ≥90% of the plasmid length. Of these 20, we found that six actually possessed *bla*_OXA-48-like_ but at a lower coverage than the threshold previously used for determining presence/absence (David et al. 2019). These findings demonstrate a strong association between presence of *bla*_OXA-48-like_ and the pOXA-48-like plasmid.

**Figure 1.**
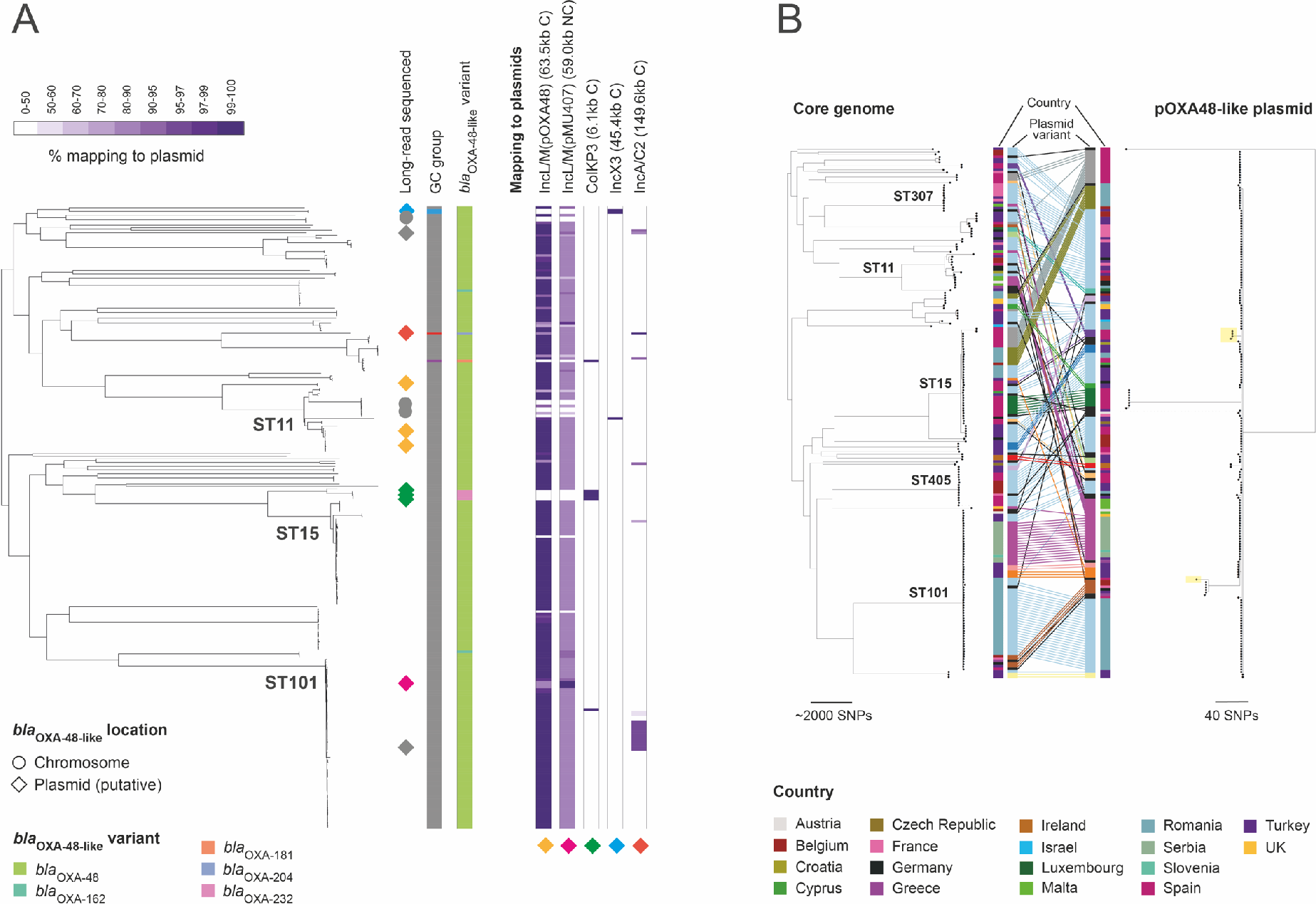
High prevalence of the pOXA-48-like plasmid sequence across *bla*_OXA-48-like_-carrying isolates. **A)** The phylogenetic tree includes 248 *bla*_OXA-48-like_-carrying isolates from *pneumoniae sensu stricto* (the single *bla*_OXA-48-like_-carrying isolate from *K. quasipneumoniae* was excluded). It was constructed using SNPs in the core genome and midpoint-rooted. All non-*bla*_OXA-48-like_-carrying isolates, which would be interspersed amongst the isolates here, were also excluded. Long-read sequenced isolates are marked next to the tree with a diamond if they carry *bla*_OXA-48-like_ on a putative plasmid sequence or a circle if they carry the gene on the chromosome. The colours of the diamonds represent distinct *bla*_OXA-48-like_-carrying plasmids that were obtained. The first two columns, from left to right, show the genetic context (GC) group of isolates assigned using the short-read assembly contigs (ambiguous isolates not assigned to any group are in grey) and the *bla*_OXA-48-like_ variant. Remaining columns show the percentage length of *bla*_OXA-48-like_-carrying plasmid sequences obtained from the hybrid assemblies that are mapped by short reads of the 248 *bla*_OXA-48-like_-carrying isolates (note the non-linear colour gradient). Mapping is shown to single representatives of the IncL/M(pOXA48) (i.e. pOXA-48-like) and ColKP3 plasmids since several highly similar plasmids were obtained with each of these replicons. The five reference plasmids used are from isolates, EuSCAPE_MT005, EuSCAPE_TR057, EuSCAPE_TR009, EuSCAPE_BE078, and EuSCAPE_FR056 (left to right in figure). Each plasmid sequence is indicated by a diamond of the same colour as that indicating the isolate(s) in the tree from which the plasmid was recovered. Mapping data for two shorter *bla*_OXA-48-like_-carrying putative plasmids is not shown (EuSCAPE_RS017 – 20.3kb; EuSCAPE_TR203 – 2.5kb). C – circular; NC – non-circular. **B)** The tanglegram links phylogenetic trees constructed using SNPs in the core genome (left) and the pOXA48-like plasmid (right). Both trees are midpoint-rooted. The trees include 207 isolates from *K. pneumoniae sensu stricto* that had mapping and bases called (A/T/C/G rather than N) at ≥90% of positions in the plasmid reference sequence. These comprise 202 isolates with *bla*_OXA-48-like_ genes and five with *bla*_VIM_ genes, the latter of which are shaded in yellow in the plasmid tree. The *bla*_OXA-48-like_-carrying isolate from *K. quasipneumoniae* that possessed this plasmid is excluded. Lines are drawn between tips in the trees representing the same isolate and coloured by the nucleotide sequence variant of the plasmid. Unique plasmid variants are coloured black. The country of origin of each isolate is shown.

Isolates carrying *bla*_OXA-48-like_ and possessing ≥99% of the IncL/M(pOXA48) plasmid sequence belonged to 37 STs across the *K. pneumoniae* species complex, and were submitted from 79 hospitals in 19 countries. These findings demonstrate the widespread nature of this plasmid. They also support a high frequency of carriage of *bla*_OXA-48-like_ by the pOXA-48-like plasmid as they rule out the possibility of a spurious association caused by lineage or geographic effects.

Despite the broad distribution of pOXA-48-like plasmids amongst chromosomal backgrounds, 122/204 (59.8%) of *bla*_OXA-48-like_-carrying isolates possessing ≥99% of this plasmid sequence belonged to one of three high-risk clonal lineages identified previously (ST11, ST15, ST101 and their derivatives) (David et al. 2019). This is approximately twice the value expected by chance (mean: 29.8%, 95% confidence intervals: 29.2-30.4%) if the distribution of pOXA-48-like plasmids mirrored the relative abundance of these clonal lineages in the sample collection (see *Methods*).

We next performed phylogenetic analysis of pOXA-48-like plasmid sequences from 202 *bla*_OXA-48-like_-carrying isolates, which included those with both mapped sequence reads and bases called (A/T/C/G rather than N) at ≥90% of reference positions. In the absence of a known outgroup, the resulting phylogenetic tree was midpoint rooted (**Figure 1B**). One hundred and seventy-six (87.1%) plasmid sequences were positioned on the ancestral node of a single main lineage or within two SNPs of this. Using published evolutionary rates for *K. pneumoniae* of 1.9 × 10^−6^ SNPs/site/year and 3.65 × 10^−6^ SNPs/site/year (Mathers et al. 2015; Stoesser et al. 2014), we estimated that the time taken for a single SNP to occur across a 63.5kb plasmid would range from 4.3 to 8.3 years. This assumed that evolutionary rates for chromosomes and plasmids are similar, which is likely given that they use the same cellular replication machinery, and are exposed to the same cellular environment. Since most pOXA-48-like sequences differ from a single ancestral variant by no more than two SNPs, this suggests that they have descended from a common ancestor that existed no more than 17 years prior to sampling (i.e. post 1996).

A tanglegram linking the plasmid-based and core genome-based phylogenies shows sharing of plasmid variants between different core genome lineages, providing clear evidence of plasmid horizontal transfer (**Figure 1B**). This has occurred frequently between core genome lineages that are co-localised at a country level. However, the core genome tree also contains 36 clonal expansions of isolates that each carry a particular plasmid variant, indicative of substantial vertical transmission. The largest contains 19 isolates from ST101, submitted from three hospitals across Romania.

Finally, all three hybrid assemblies harbouring the *bla*_OXA-48_ variant in the chromosome carried the gene within a Tn*6237* composite transposon, which is a ~20kb sequence that also carries *bla*_OXA-48-like_ in the pOXA-48-like plasmids (**Supplementary Figure 2**). We found evidence of at least two independent chromosomal integrations of Tn*6237* in ST11 and ST530, respectively, and subsequent clonal spread (see **Supplementary Note**).

### Spread of *bla*_VIM_ and *bla*_NDM_ genes mediated by transient associations of diverse plasmids with multiple lineages

We obtained hybrid assemblies carrying *bla*_VIM_ genes representing the ten GC groups identified (**Supplementary Table 3**). Amongst these, we found *bla*_VIM_ in putative plasmid sequences (46.0-284.3kb) in 9/10 hybrid assemblies and in one chromosomal sequence (5.3Mb). Another putative plasmid sequence harbouring *bla*_VIM_ was obtained from an isolate harbouring two carbapenemase genes (carrying also *bla*_KPC_) but excluded from further analyses due to the short contig length (2.9kb). We also obtained hybrid assemblies carrying *bla*_NDM_ genes representing the 15 GC groups identified (**Supplementary Table 3**). All carried *bla*_NDM_ on putative plasmid sequences (12.2-197.6kb). Overall, *bla*_VIM_ and *bla*_NDM_-carrying plasmids harboured diverse replicon types. Several also shared partial structural and/or sequence homology.

The same short-read mapping approach used previously allowed us to determine the prevalence of the different plasmid sequences across all *bla*_VIM_ (*n*=56) and *bla*_NDM_-carrying (*n*=79) isolates in the sample collection. Two of the *bla*_VIM_-carrying plasmids (from EuSCAPE_LV006 and EuSCAPE_IT312) and one of the *bla*_NDM_-carrying plasmids (from EuSCAPE_RS064) were mapped over ≥90% by only the long-read sequenced isolate and thus were unique in this collection (**Figure 2**). However, many plasmids were associated with clonal expansions of isolates, which were defined as two or more same-ST isolates clustered in the core genome-based phylogeny. In particular, 39/56 (69.6%) *bla*_VIM_-carrying isolates belonged to six clonal expansions and 38/79 (48.1%) *bla*_NDM_-carrying isolates belonged to seven clonal expansions, each with ≥99% mapping to a particular plasmid. Isolates from ST11, ST15 and ST101 (and their derivatives) accounted for the majority (56.4% and 71.1%, respectively) of these.

**Figure 2.**
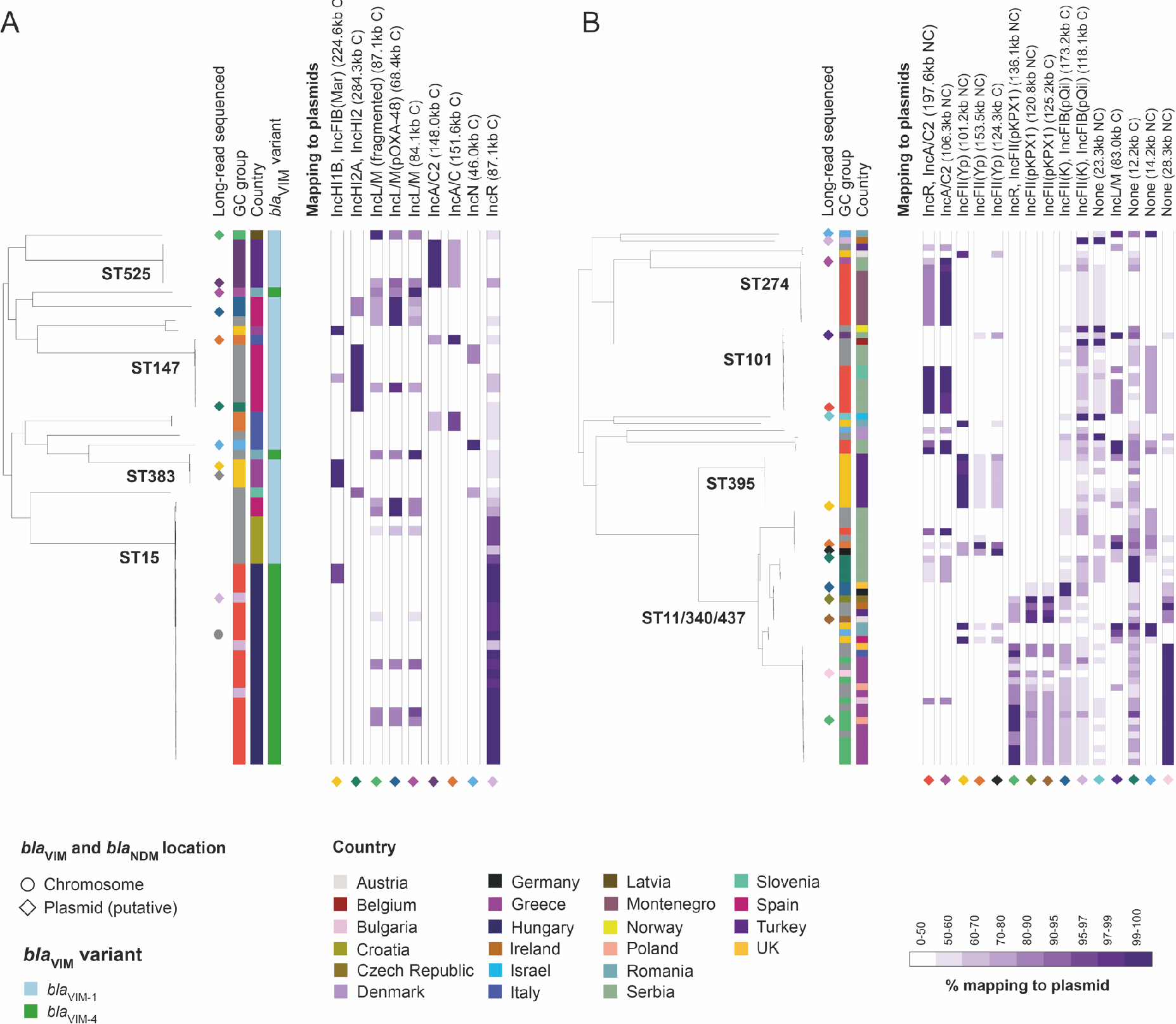
Plasmids carrying *bla*_VIM_ and *bla*_NDM_ genes are associated with individual clonal expansions. The phylogenetic trees, constructed using SNPs in the core genome, show 56 *bla*_VIM_-carrying isolates **(A)** and 79 *bla*_NDM_-carrying isolates **(B)** from *K. pneumoniae sensu stricto*. Both trees are midpoint-rooted. We excluded all non-*bla*_VIM_ and non-*bla*_NDM_-carrying isolates, respectively, which would be interspersed amongst the isolates here. Long-read sequenced isolates are marked next to the tree with a diamond if they carry the carbapenemase gene on a putative plasmid sequence or a circle if they carry the gene on the chromosome. The colours of the diamonds represent distinct carbapenemase-carrying plasmids that were obtained. Columns, from left to right, show the genetic context (GC) group of isolates assigned using the short-read assembly contigs (ambiguous isolates not assigned to any group are in grey), the country of isolation and the gene variant (for *bla*_VIM_ genes only as all *bla*_NDM_ genes were *bla*_NDM-1_). Remaining columns show the percentage length of putative plasmids carrying *bla*_VIM_ (A) and *bla*_NDM_ (B) genes obtained from the hybrid assemblies that were mapped by short reads (note the non-linear colour gradient). The nine reference plasmids used in (A) are from isolates, EuSCAPE_GR073, EuSCAPE_ES094, EuSCAPE_LV006, EuSCAPE_ES220, EuSCAPE_RO094, EuSCAPE_TR203, EuSCAPE_IT062, EuSCAPE_IT312, EuSCAPE_HU009 (left to right in figure). The fifteen used in (B) are from EuSCAPE_RS017, EuSCAPE_RS105, EuSCAPE_TR083, EuSCAPE_RS064, EuSCAPE_RS002, EuSCAPE_PL046, EuSCAPE_CZ007, EuSCAPE_AT023, EuSCAPE_DE019, EuSCAPE_IE008, EuSCAPE_IL075, EuSCAPE_RS010, EuSCAPE_RS081, EuSCAPE_RO052 and EuSCAPE_GR094 (left to right in figure). Each putative plasmid sequence is indicated by a diamond with the same colour as that indicating the isolate in the tree from which the plasmid was recovered. Mapping data for one shorter *bla*_VIM_-carrying putative plasmid is not shown (EuSCAPE_GR075 - 2.9kb). C – circular; NC – non-circular.

The core genome diversity within clonal expansions associated with particular plasmids was typically low with maximum pairwise SNP differences per clonal expansion ranging from 0-51 (median, 8) for *bla*_VIM_-carrying isolates and 8-54 (median, 17) for *bla*_NDM_-carrying isolates. Given the published mutation rates (Mathers et al. 2015; Stoesser et al. 2014), we estimated the time taken for two isolates to diverge from a common ancestor by 54 SNPs would be 3.0 to 5.8 years. This is suggestive of recent acquisition of the plasmids by these lineages, and indicates that associations between the chromosome and these plasmids may be often only transient. Indeed, 4/6 and 2/7 clonal expansions of *bla*_VIM_-carrying and *bla*_NDM_-carrying isolates were restricted to a single hospital (and correspondingly have few SNP differences). A further 2/6 and 1/7, respectively, contained isolates submitted from different hospitals in the same country, while 4/7 of those with *bla*_NDM_-carrying isolates were from different countries. While isolates from the three high-risk clonal lineages constituted 7/13 of the total clonal expansions associated with particular *bla*_VIM_ and *bla*_NDM_ plasmids, they accounted for 6/7 of those that had spread to multiple hospitals or countries.

We found some indications of plasmid sharing between STs, with three circularised *bla*_VIM_-carrying plasmids (from EuSCAPE_GR073, EuSCAPE_ES220 and EuSCAPE_RO094) and two circularised *bla*_NDM_-carrying plasmids (from EuSCAPE_IE008 and EuSCAPE_RS010) mapped with short reads across ≥99% of their length by isolates from different STs (**Figure 2**). These often included isolates from different STs submitted from the same country but never the same hospital. Notably, they included a 68.4kb *bla*_VIM_-1-carrying plasmid with high similarity to the pOXA-48-like plasmids that was recovered from the hybrid assembly of a ST483 isolate (EuSCAPE_ES220) but found also in ST11 and ST15 using short-read mapping (**Figure 1B** and **Supplementary Figure 3**; see **Supplementary Note**).

While plasmids shared between lineages were in general observed rarely, we found one exception, which was a non-circularised *bla*_NDM_-carrying IncA/C2 plasmid sequence recovered from a ST274 isolate (EuSCAPE_RS105). This was mapped across ≥99% of its length by another eight ST274 isolates but also four ST101, two ST147 and one ST437 isolate(s). Long-read sequencing of one of the ST101 isolates (EuSCAPE_RS017) demonstrated that this IncA/C2 sequence formed part of a larger plasmid sequence in this isolate comprising both IncA/C2 and IncR replicons. We could not find the additional plasmid sequence, nor evidence of an IncR replicon, within the assembly of EuSCAPE_RS105. However, only three SNPs were found across 101.4kb of shared sequence between the two plasmids (**Supplementary Figure 4)**, suggestive of recent common ancestry.

### Dissemination of *bla*_KPC_ genes by stable association with ST258/512 despite frequent mobilisation between diverse plasmids

We obtained 44 *bla*_KPC_-carrying hybrid assemblies from isolates representing 43 GC groups. These included two isolates from the same group selected to investigate possible plasmid transfer between STs (for details on these, see **Supplementary Note; Supplementary Figure 5)**. The *bla*_KPC_ gene was found on a chromosomal sequence (3.8Mb) in one hybrid assembly, and on putative plasmid sequences (7.9-313.6kb) in the remaining 43. We found diverse replicon types amongst these putative plasmids, including those from the single clonal lineage of ST258/512. This lineage contains 230/312 (73.7%) of all *bla*_KPC_-carrying isolates in the sample collection.

Pairwise sequence comparisons between 24 circularised *bla*_KPC_-carrying plasmids indicated that 15 were structural variants of two major IncF backbone types: backbone I (*n*=9) and backbone II (*n*=6) (**Figure 3**). The first backbone type represents pKpQIL-like plasmids and we found that two have an identical size and structure to the originally described pKpQIL plasmid (Leavitt et al. 2010) (**Supplementary Figure 6**). The second backbone type shares sequence with pKPN3 (accession number, CP000648) but also pKpQIL-like (backbone I) plasmids (**Supplementary Figure 7**). Both backbone types I and II were found in ST258/512 and were geographically dispersed. Overall we found poor concordance between the plasmid types carrying *bla*_KPC_ genes and the phylogeny of the host strain, including within ST258/512 (**Figure 3)**.

**Figure 3.**
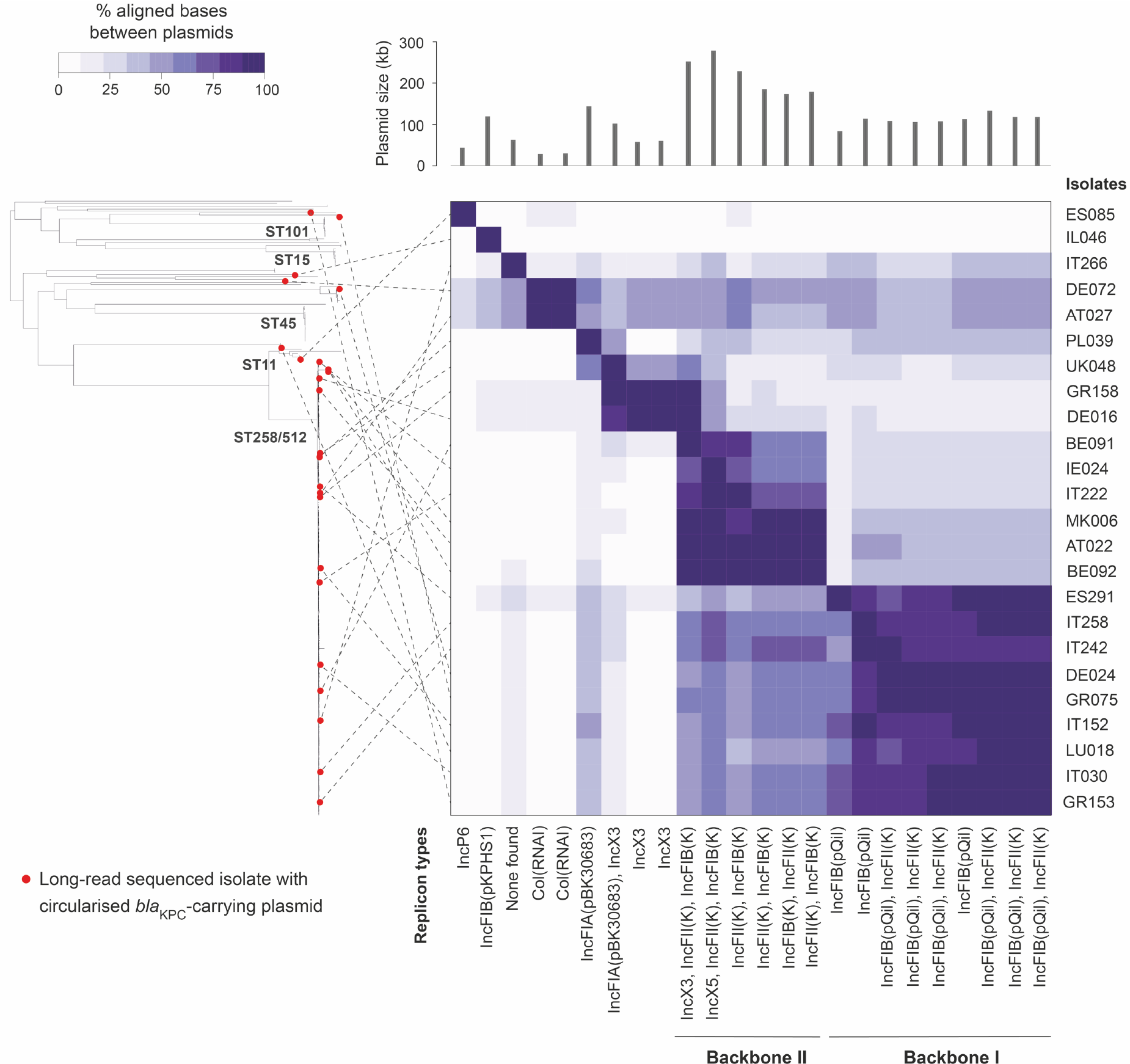
Comparison of 24 circularised *bla*_KPC_-carrying plasmids shows dominance of two major IncF backbone types. The phylogenetic tree contains 311 *bla*_KPC_-carrying isolates from *K. pneumoniae sensu stricto* (the single *bla*_KPC_-carrying isolate from *K. variicola* is excluded). The tree was constructed using SNPs in the core genome and is midpoint-rooted. Twenty-four isolates from which circularised *bla*_KPC_-carrying plasmids were obtained are marked by red circles in the tree. The heat map shows the percentage of bases in each plasmid that could be aligned to each of the other plasmids using NUCmer (the row and column orders are the same). Dotted lines link the 24 long-read sequenced isolates in the phylogenetic tree to their respective plasmids in the heat map. The plasmid length and replicon types found in each plasmid are shown above and below the heat map, respectively.

Short read mapping of all *bla*_KPC_-carrying isolates in the sample collection (*n*=312) to the newly-obtained *bla*_KPC_-carrying plasmids demonstrated that some (i.e. the IncP6, IncN and IncFIB(pKPHS1) plasmids) were found only rarely (**Supplementary Figure 8**). However, many isolates carry two or more of the reference plasmids, including 66 ST258/512 isolates that have ≥99% mapping to four distinct plasmid types with ColRNAI, IncX3, IncFII(K)/IncFIB(pQIL) (i.e. backbone I) and IncFII(K)/IncFIB(K) (i.e. backbone II) replicons. This means that several of the reference plasmid sequences are frequently present in the same isolate, either in the same or a different structural arrangement, and we cannot infer which one (or more) contains the carbapenemase gene using the short sequence reads.

We therefore used an alternative approach that takes advantage of *bla*_KPC_ genes typically being on longer contigs in the short-read genome assemblies than other carbapenemases (**Table 1**). We compared each of the short-read contigs harbouring *bla*_KPC_ genes (*n*=312) with each of the 24 circularised *bla*_KPC_-carrying plasmids from the hybrid assemblies. If ≥98% of the contig sequence could be aligned to a plasmid, we considered this as a match (**Supplementary Figure 9**; see **Supplementary Note**). We found that 28/82 (34.1%) of short-read contigs from non-ST258/512 isolates matched either backbone I or II plasmids. This contrasted with 200/230 (87.0%) of contigs from ST258/512 isolates. Of these 200, 183 (79.6% of 230) were not compatible with any other plasmid types. These results support backbones I and II (or related variants of these) being the dominant vectors of *bla*_KPC_ genes in ST258/512. However, only 36/230 (15.7%) and 28/230 (12.2%) ST258/512 contigs could be unambiguously assigned to either backbone I or II, respectively.

We next aimed to understand the evolutionary processes that have led to *bla*_KPC_ genes being carried on diverse plasmids within ST258/512, with a low degree of congruence between the plasmid type carrying *bla*_KPC_ and the core genome-based phylogeny. First, we determined if the plasmids on which we found *bla*_KPC_ are stably associated with the ST258/512 lineage or have been acquired repeatedly from outside of the lineage. We constructed phylogenetic trees of 91 pKpQIL-like plasmids and 135 IncX3 plasmids from ST258/512 isolates. These were from isolates that had short reads that mapped to ≥99% of the plasmid reference sequences, and comprised 48.9% and 95.1% of ST258/512 isolates possessing a IncFIB(pQIL) and IncX3 replicon, respectively. Comparisons of these plasmid-based trees with a core genome-based phylogeny of ST258/512 isolates revealed shared evolutionary histories, suggestive of single acquisitions early in the lineage history (**Figure 4**). IncX3 plasmid sequences with and without a *bla*_KPC_ gene (as confirmed using the hybrid assemblies) were also intermingled in the phylogenetic tree of IncX3 plasmids, further indicative of vertical propagation of the plasmid within the lineage coupled with occasional gain and/or loss of *bla*_KPC_ (**Figure 4B**). While phylogenetic reconstructions were not undertaken for the ColRNAI plasmid (due to a lack of diversity) or the backbone II plasmid (due to very high gene content variation), sequence comparisons of these plasmids with and without *bla*_KPC_ genes showed high similarity between their backbones, indicative of a common origin (**Supplementary Figures 10** and **11;** see **Supplementary Note**). These findings suggest that *bla*_KPC_ genes can be maintained by plasmid backbones that are stable within the lineage. However, they do not reveal whether *bla*_KPC_ genes have moved between plasmids within a single cell, or whether they have repeatedly been imported into ST258/512 plasmids from other strains (from either within or outside of ST258/512).

**Figure 4.**
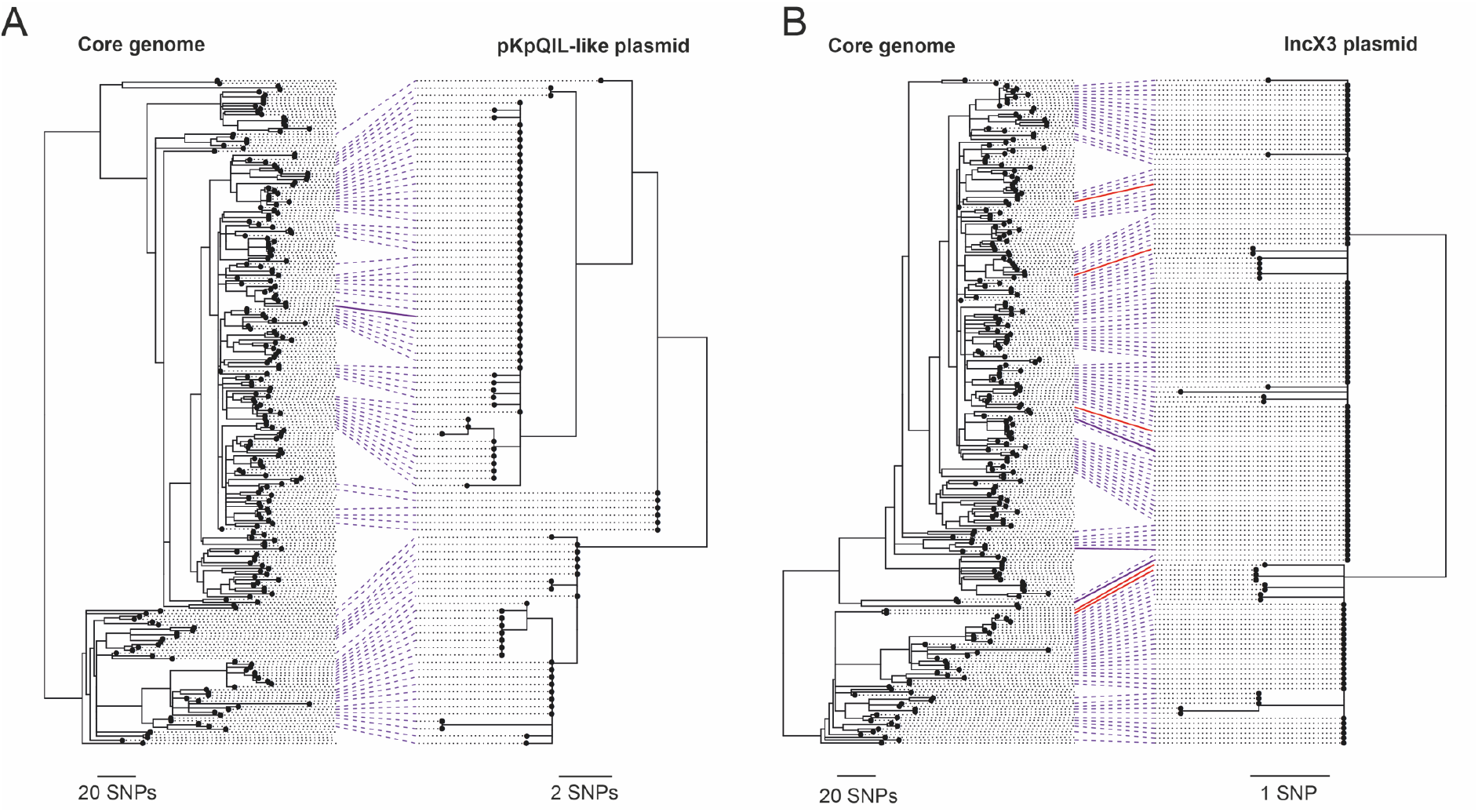
High congruence between pKpQIL-like and IncX3 plasmid phylogenies with the core genome phylogeny of ST258/512 reveals shared evolutionary histories. Each tanglegram comprises a phylogeny of the ST258/512 lineage constructed using all SNPs in the core genome (mapping-based) alignment and either the pKpQIL-like **(A)** or IncX3 **(B)** plasmids. The core genome phylogenies include all 236 ST258/512 isolates and were rooted based on previous phylogenetic analyses of the full sample collection that included outgroups of ST258/512 (David et al. 2019). Ninety-one pKpQIL-like and 135 IncX3 plasmid sequences from isolates that had bases (A/T/C/G) called at ≥99% positions in the plasmid reference were included in the plasmid phylogenies. These were rooted to provide the highest concordance with the core genome phylogenies. Lines are drawn between tips in the two trees representing the same isolate. The solid purple lines indicate isolates which were found to carry *bla*_KPC_ on a pKpQIL-like (A) or IncX3 (B) plasmid in the hybrid assemblies. Red lines (in (B) only) indicate isolates that were found to carry *bla*_KPC_ on an alternative plasmid to an IncX3 plasmid in the hybrid assemblies.

To distinguish between these possibilities, we next used the TETyper tool (Sheppard et al. 2018), which takes short-read data as input, to screen all *bla*_KPC_-carrying isolates for the ~10kb Tn*4401* transposon and investigate its patterns of inheritance. This transposon is known from previous studies to be the major carrier of *bla*_KPC_ genes in *K. pneumoniae*, especially amongst European strains (Cuzon et al. 2011; Chen et al. 2014). Tn*4401* sequences were found in 229/230 (99.6%) ST258/512 isolates harbouring *bla*_KPC_ genes, and classified into “combined” variants based on both structural and SNP variation. We found two predominant combined variants, Tn*4401*a-1 (*n*=42) and Tn*4401*a-2 (*n*=176), which differ by a single SNP that also distinguishes the *bla*_KPC_-2 and *bla*_KPC_-3 gene variants. These variants correlate well with the core genome-based phylogeny of ST258/512, with 42/46 (91.3%) isolates in one major clade (clade 1) carrying Tn*4401*a-1, and 175/184 (95.1%) isolates in the second major clade (clade 2) carrying Tn*4401*a-2 (**Figure 5**). This indicates a single major acquisition of Tn*4401* (carrying *bla*_KPC_) by an ancestor of this lineage, followed by relatively stable association during the clonal expansion of ST258/512. Taken together, the combined stability of both the plasmids and Tn*4401* transposon within the ST258/512 lineage suggests that Tn*4401* (carrying *bla*_KPC_) has moved primarily between plasmids in the same bacterial cell or between genetically identical cells (such as those in a clonal infection). We also cannot rule out movement of Tn*4401* between plasmids from different strains, provided that these strains are from the same major clade of ST258/512 (i.e. possess the same combined variant of Tn*4401*). However, the data is not compatible with frequent movement of Tn*4401* between strains from different major clades, or frequent import of Tn*4401* into ST258/512 from outside of the lineage.

**Figure 5.**
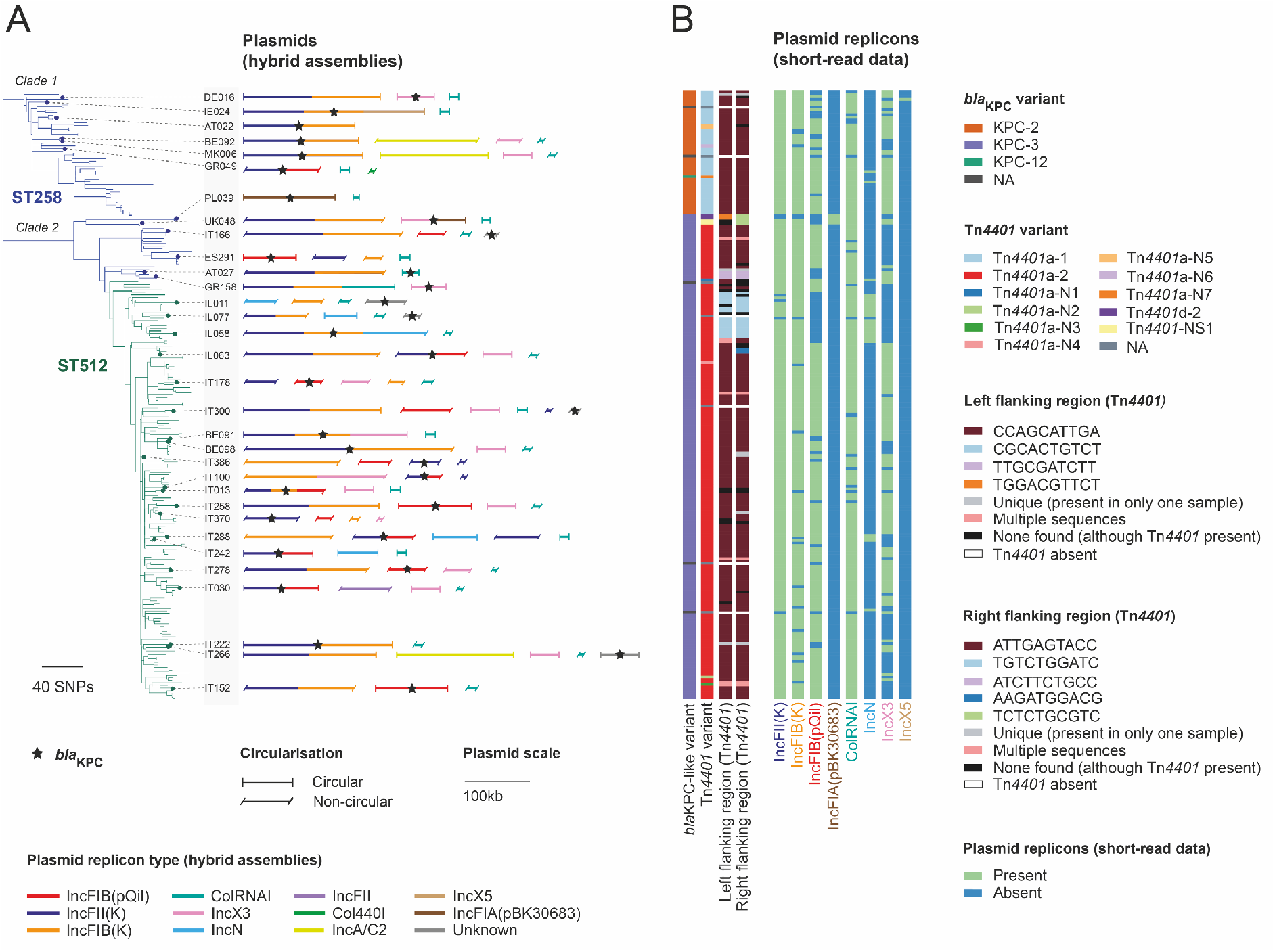
Movement of *bla*_KPC_ genes between plasmids in the ST258/512 lineage. **A)** The phylogenetic tree contains 236 isolates belonging to ST258/512. It was constructed using SNPs from a core genome (mapping-based) alignment and rooted based on previous phylogenetic analyses of the full sample collection that included outgroups of ST258/512 (David et al. 2019). Thirty-two long-read sequenced isolates carrying *bla*_KPC_ on a putative plasmid sequence are indicated by small circles on the tree tips. Putative plasmid sequences derived from the hybrid genome assemblies with at least one known replicon type and/or containing *bla*_KPC_ are depicted next to the tree. These are scaled by size and coloured by any replicon types found in the sequence. A star indicates the presence of *bla*_KPC_ within these sequences. **B)** Metadata columns, from left to right, show the *bla*_KPC_ variant, the Tn*4401* variant, the 10bp left and right flanking regions of Tn*4401*, and presence or absence of eight plasmid replicon types found using short-read data to be associated with *bla*_KPC_ in the hybrid assemblies. Tn*4401a*-N1 – Tn*4401a*-N7 represent novel SNP variants of the structural variant, Tn*4401a*. Tn*4401*-NS1 represents a novel structural variant of Tn*4401*.

Finally, we investigated whether Tn*4401* (and *bla*_KPC_) has moved between plasmids via transposition or as part of larger recombination events. Using the short-read data with TETyper, we found identical 10bp flanking regions (CCAGCATTGA/ATTGAGTACC) upstream and downstream of Tn*4401* in 176/230 (76.5%) ST258/512 isolates, which include 5bp ATTGA target site duplications (**Figure 5**). Amongst *bla*_KPC_-carrying plasmids from the hybrid assemblies, these particular flanking regions were restricted to backbone I and II plasmids (and three putative plasmid sequences with no known replicons). Taking these flanking regions to be markers of backbone I and II plasmids, these findings further support the dominant role of the two backbone types as vectors of *bla*_KPC_ genes. They also indicate that *bla*_KPC_ genes are typically mobilised between these plasmid types by larger recombination events (i.e. >10kb) which transfer Tn*4401* together with additional flanking sequence. Indeed we found a shared 34kb sequence region around *bla*_KPC_ in plasmids representing backbones I and II that were recovered from closely-related isolates (EuSCAPE_GR049 and EuSCAPE_MK006) (**Supplementary Figure 12**). Only two SNPs were found across this region, suggestive of recent transfer, in contrast to several hundred SNPs found across the remaining homologous sequence. Conversely, we found distinct Tn*4401* flanking regions in the ColRNAI, IncX3, and IncFIA(pKB30683) plasmids carrying *bla*_KPC_ genes in the hybrid assemblies, indicative of transposition of Tn*4401* into these plasmids (**Figure 5**). We also identified two different flanking regions both upstream and downstream of Tn*4401* in five ST258/512 isolates, suggesting that *bla*_KPC_ is present in two copies. However, none of these five isolates were long-read sequenced to verify this.

## Discussion

Molecular and genomic surveillance systems for bacterial pathogens currently rely on tracking clonally-evolving lineages. By contrast, extra-chromosomal plasmids, which can spread horizontally between strains and even species (Mathers et al. 2011; Martin et al. 2017), are usually excluded or analysed with low-resolution techniques (such as Inc typing). This is despite plasmids being the primary carriers of antibiotic resistance genes across many key pathogens. Here, we used combined long- and short- read sequencing of isolates from a European structured survey (EuSCAPE) (Grundmann et al. 2017; David et al. 2019) to investigate the diversity, distribution and transmission dynamics of resistance plasmids in *K. pneumoniae.* We focused on plasmids carrying carbapenemase genes, which confer resistance to carbapenems, a last-line class of antibiotics. We identified three major patterns by which carbapenemase genes have disseminated via plasmids, summarised as using one plasmid/multiple lineages (*bla*_OXA-48-like_), multiple plasmids/multiple lineages (*bla*_VIM_ and *bla*_NDM_) and multiple plasmids/one lineage (*bla*_KPC_). Despite these contrasts, our work revealed the high dependency of all three modes of carbapenemase gene spread on a small number of high-risk clones.

Previous studies have demonstrated a dominance of high-risk clones among antibiotic-resistant *K. pneumoniae* infections (Munoz-Price et al. 2013), although the reasons driving their success are still debated (Mathers et al. 2015). Here we have shown that carbapenemase-carrying plasmids are acquired by diverse lineages, such as the pOXA-48-like plasmid that was found in 37 STs. Yet our phylogenetic analyses indicate that carbapenemase-carrying plasmids are (i) non-randomly associated with high-risk clones (i.e. ST11, ST15, ST101, ST258/512), (ii) propagating by clonal expansion, and (iii) frequently spreading across healthcare networks and national borders. These findings reinforce the importance of preventing transmission, particularly of high-risk STs, through early detection and rigorous infection control.

The first *bla*_OXA-48_-carrying isolate described in 2004 (Poirel et al. 2004) was later found to carry the carbapenemase gene within a IncL/M pOXA-48-like plasmid (Poirel et al. 2012). Since then, numerous studies have reported this plasmid as the dominant vector of *bla*_OXA-48-like_ genes both within and outside of Europe, and in both *K. pneumoniae* and other *Enterobacterales* species (Skalova et al. 2017; Potron et al. 2013). It has also since been further distinguished as an IncL plasmid, after IncL and IncM plasmids were found to be genetically distinct and compatible (Carattoli et al. 2015). Experiments performed by Adler et al. (2016) demonstrated that pOXA-48-like plasmids show very efficient conjugation both within and between bacterial species, helping to explain their predominance. Here, our comparative analyses demonstrate a single main acquisition of a *bla*_OXA-48-like_ gene by a pOXA-48-like backbone. They suggest that almost all pOXA-48-like plasmids in our collection share a common ancestor that existed fewer than 17 years prior to the sample collection (i.e. post 1996). These plasmids have since spread rapidly as we found them in 79 hospitals in 19 countries across Europe. Most notably, we have shown that, despite frequent horizontal transfer of the plasmid, this onward spread has been primarily driven by the clonal expansion of high-risk STs.

By contrast, we found *bla*_VIM_ and *bla*_NDM_ genes on multiple diverse plasmids, which is concordant with reports from the literature (Samuelsen et al. 2011; Perez-Vazquez et al. 2019). As with the pOXA-48-like plasmid, our results show that clonal expansions, especially of high-risk STs, have driven the spread of these plasmids. We noted that associations of clonal lineages with plasmids carrying *bla*_VIM_ and *bla*_NDM_ genes were mostly recent, suggesting that they may be typically only transient. While the maximum age of any clonal expansion associated with a particular plasmid was estimated to be 5.8 years, most were much younger than this. We also found that plasmid sharing between lineages was coupled with structural changes in the plasmids that accumulate over time. We propose that high rates of recombination and rearrangement amongst plasmids could partially explain both the transient associations between lineages and plasmids, as well as the absence of any single dominant plasmid found across multiple lineages.

The most striking example of the reliance on high-risk clones is provided by the *bla*_KPC_ gene. Since its discovery in 1996, this gene has disseminated worldwide at a remarkable pace (Munoz-Price et al. 2013). A single clonal lineage, ST258/512, accounted for >70% of all *bla*_KPC_ genes found amongst the EuSCAPE sample collection (David et al. 2019). Previous studies have shown that *bla*_KPC_ can be carried by different plasmids in ST258/512 (Pitout et al. 2015; Noll et al. 2018). In particular, the pKpQIL-like (backbone I) plasmids have been highlighted as important vectors in North America, Europe and the Middle East (Chen et al. 2014; Papagiannitsis et al. 2016; Doumith et al. 2017). However, the origin of the different *bla*_KPC_-carrying plasmids was unknown. Here we have shown that several of the key plasmid types carrying *bla*_KPC_ genes, including pKpQIL-like plasmids, are stably associated with the ST258/512 lineage. Our data support a single acquisition of *bla*_KPC_ by an early ancestor of the lineage, followed by movement of the gene between different plasmid types in the same bacterial cell. This was coupled with frequent recombination and rearrangement events between different plasmid types, leading to a complex array of mosaic structures carrying *bla*_KPC_ genes in the ST258/512 lineage.

We acknowledge several limitations of our study. First, we required carbapenemase-carrying contigs in the short-read assemblies to have ≥4 genes to be used for defining GC groups, which then guided selection of isolates for long-read sequencing. This may have reduced the amount of plasmid diversity captured by disregarding isolates carrying carbapenemase genes within particular repetitive structures. Second, we used only one isolate from each GC group for long-read sequencing. This meant we were unable to assess the structural diversity and evolution of plasmids within shorter timescales, such as within clonal expansions of *bla*_VIM_ and *bla*_NDM_-carrying isolates. We could also not confirm the stable presence of the carbapenemase gene on particular plasmids within these clonal expansions. Finally, the use of short-read mapping to reference plasmids had varying levels of utility and appropriateness. While it was useful for identification and phylogenetic analyses of stable plasmids (e.g. the pOXA-48-like plasmid), the common occurrence of mosaic plasmids could make these data difficult to interpret.

In summary, we have highlighted three major modes of carbapenemase gene spread via plasmids. Consideration of each will be vital for incorporation of high-resolution plasmid data into more comprehensive genomic surveillance systems.

## Materials & Methods

### Clustering of short-read assembly contigs carrying carbapenemase genes into genetic context (GC) groups

Carbapenemase genes (*bla*_OXA-48-like_, *bla*_VIM_, *bla*_KPC_ and *bla*_NDM_) were detected in the previously generated short-read assemblies (David et al. 2019) using BLASTn (Altschul et al. 1990). A minority of carbapenemase genes were not found in the assemblies despite detection using raw sequence reads. Conversely, carbapenemase genes that were detected in the assemblies but found with low coverage using the raw sequence reads (<0.2x the coverage of the MLST gene with the lowest coverage) were excluded. All remaining contigs carrying carbapenemase genes were extracted from the short-read assemblies and annotated using Prokka v1.5 (Seemann, 2014). Contigs with four or more genes (including the carbapenemase) were used in the subsequent clustering analysis.

Annotated contigs containing a particular carbapenemase gene were used as input to Roary v3.11.3 (Page et al. 2015) to cluster the genes from different contigs into groups based on their nucleotide identity. Roary was run using default settings with the exception of the addition of the “-s” flag to prevent genes that are presumed to be paralogous being split into different gene groups. Contigs were excluded if the carbapenemase gene lacked a proper start or stop codon, as detected by Roary.

The remaining contigs were assigned to clustering groups based on the order and groupings of genes surrounding the carbapenemase (**Table S1**). Those with the same genes (according to the Roary gene groupings) in the same order around the carbapenemase were assigned to the same clustering group, while those with different genes and/or a different order were separated into different clustering groups. Contigs assigned to the same clustering group could be of different lengths and have different numbers of genes, with some contigs extending beyond others. Contigs could also belong to multiple clustering groups (i.e. if they matched multiple, usually larger contigs that are themselves different). Contigs that belonged to a single clustering group were assigned to a genetic context (GC) group, while those that belonged to multiple clustering groups were identified as having an “ambiguous” genetic context (**Table S2**). This clustering method is equivalent to finding all maximal cliques in a graph (Eppstein et al. 2013) and the solutions were obtained using a C++ script (https://github.com/darrenstrash/quick-cliques).

### Culture, DNA extraction and long-read sequencing

Seventy-nine isolates were selected for long-read sequencing (**Table S3**). These include one isolate from each of the GC groups, with the exception of one *bla*_OXA-48-like_ group and two *bla*_KPC_ groups for which representative isolates were unavailable. An additional eight *bla*_OXA-48-like_-carrying isolates were also included, while two *bla*_KPC_-carrying isolates from the same GC group but different STs were selected to investigate possible within-hospital plasmid transfer.

The samples were grown on MacConkey agar plates at 37°C overnight and this was repeated until single colonies were visible on the plates. Single colonies of each sample were grown overnight in 7mL of low salt lysogeny broth (LB) in a shaking incubator. Three mL of this culture was spun down at the maximum number of revolutions per minute (RPM) for three minutes and the supernatant was discarded. The pellet was resuspended in 500μL of lysis buffer, incubated at 80°C for five minutes and then cooled. Two hundred and seventy-five μL of 3M sodium acetate was added to the samples, which were vortexed briefly to mix. They were then spun at the maximum RPM for ten minutes. The supernatant was removed and added to clean Eppendorf tubes. Five μL of RNAse A was added and the samples incubated for 30 minutes at 37°C before cooling. Two hundred μL of protein precipitate solution (Promega) was added, and the samples were incubated on ice for five minutes before being spun at the maximum RPM for three minutes. Finally, ethanol precipitation was performed.

Library preparation for all isolates was performed using the SMRTbell Template Prep Kit 1.0. Long-read sequencing of 39 isolates was performed on the RSII instrument from Pacific Biosciences using C4/P6 chemistry, and 40 isolates were sequenced on the Sequel instrument using v2.1 chemistry and a multiplexed sample preparation. The Sequel data was demultiplexed using Lima in the SMRT link software (https://github.com/PacificBiosciences/barcoding).

### Hybrid assembly

The long-read data were assembled together with the previously obtained short-reads for each sample (David et al. 2019) using the hybrid assembler, Unicycler v0.4.7 (Wick et al. 2017). Default settings were used for all samples with the exception of four (EuSCAPE_IL075, EuSCAPE_DE024, EuSCAPE_ES089, EuSCAPE_TR203). For these, we set the flag “--depth_filter”, which represents the fraction of the chromosomal depth below which contigs are filtered out, to 0.1 since the carbapenemase genes were absent from the assemblies when the default setting of 0.25 was used. Assembly statistics were generated using QUAST v.4.6.0 (Gurevich et al. 2013). Assemblies were annotated using Prokka v1.5 (Seemann, 2014). Annotated assemblies are available in the European Nucleotide Archive (ENA) and accession numbers can be found in **Table S3**.

### Characterisation of hybrid assemblies

Contigs containing the carbapenemase genes were identified in the hybrid assemblies using BLASTn (Altschul et al. 1990). Replicon typing of all contigs was performed using Ariba v2.6.1 (Hunt et al. 2017) with the PlasmidFinder database (Carattoli et al. 2014). Galileo AMR (Partridge & Tsafnat, 2018) was used to annotate mobile genetic elements within carbapenemase-carrying plasmids.

### Plasmid comparisons

NUCmer v3.1 from the MUMmer package (Kurtz et al. 2004) was used to determine the length of sequence that could be aligned between pairs of plasmids, and the number of single nucleotide polymorphisms (SNPs) amongst the aligned regions. The Artemis Comparison Tool (ACT) v13.0.0 (Carver et al. 2005) was used to compare and visualise structural variation between two or more sequences.

### Short-read mapping to plasmid sequences

Mapping of the previously generated short sequence reads (David et al. 2019) to putative plasmid sequences obtained from the hybrid assemblies was used to determine the length of the reference plasmid sequence present across isolates. Sequence reads were mapped using Burrows Wheeler Aligner (Li & Durbin, 2009) and an in-house pipeline was used to identify SNPs using SAMtools mpileup v.0.1.19 and BCFtools v1.2 (Li et al. 2009). The length of the reference plasmid that was mapped by a minimum of one sequence read in each sample was determined from the BAM file. Upon testing of this approach, we found that the short reads of each carbapenemase-carrying isolate from which we obtained a hybrid assembly mapped to 99.8-100% (median, 100%) of the putative carbapenemase-carrying plasmid sequence from the hybrid assembly.

### Core genome-based phylogenetic analyses

Phylogenetic trees comprising 248 *bla*_OXA-48-like_, 56 *bla*_VIM_, 311 *bla*_KPC_ and 79 *bla*_NDM_-carrying isolates of *K. pneumoniae sensu stricto* were each constructed using variable positions within an alignment of 2539 genes. These represent loci that were found to be “core” genes (i.e. present in at least 95% of isolates within each species of the *K. pneumoniae* species complex) in previous analyses (David et al. 2019). Phylogenetic trees were inferred using RAxML v8.2.8 (Stamatakis, 2006) and midpoint-rooted. The same core genome alignment was also used to calculate pairwise SNP differences between isolates.

The phylogenetic tree of the ST258/512 lineage was constructed by mapping the short reads of the isolates to an ST258 reference genome (NJST258_1 (Deleo et al. 2014)). Recombined regions were removed from the pseudo-genome alignment using Gubbins v1.4.10 (Croucher et al. 2015) and the variable sites in the resulting alignment were used as input to RAxML v8.2.8 (Stamatakis, 2006). The tree was rooted based on previous phylogenetic analyses of the full sample collection that included outgroups of ST258/512 (David et al. 2019).

### Plasmid-based phylogenetic analyses

To construct a phylogenetic tree of pOXA-48-like plasmids, we first generated a plasmid alignment comprising *bla*_OXA-48-like_-carrying isolates with bases mapped and called at ≥90% of the reference plasmid from EuSCAPE_MT005. This included plasmids from 203 isolates, although one plasmid sequence from the *K. quasipneumoniae* species was subsequently excluded. An additional five *bla*_VIM_-carrying isolates with bases mapped and called at ≥90% of the reference plasmid were also included in the alignment.

Phylogenetic trees of the pKpQIL-like and IncX3 plasmids from the ST258/512 lineage were also constructed using the alignments generated from mapping of the short reads to reference plasmids. The references used were the pKpQIL-like plasmid of EuSCAPE_IT030 and the IncX3 plasmid of EuSCAPE_IL063. Isolates with ≥99% of bases called at the reference positions were included.

Variable positions in these plasmid alignments, excluding any positions containing an N (rather than A/T/C/G) in ≥1 isolate, were used to infer phylogenetic trees using RAxML v8.2.8 (Stamatakis, 2006). Nucleotide variants of plasmids were determined using the same alignments. Tanglegrams linking the core genome and plasmid phylogenies were generated using the “cophylo” function from the “phytools” package in R (https://www.r-project.org/).

### Comparison of the actual and expected proportion of pOXA-48-like plasmids in high-risk lineages

We compared the actual proportion of pOXA-48-like plasmids carried amongst three high-risk clonal lineages (ST11, ST15, ST101) with the expected proportion if the distribution of plasmids reflected the relative abundance of these lineages in the population. Isolates belonging to other STs that have evolved from these three STs were included with them, with the exception of ST258/512. We first determined whether each isolate in the full EuSCAPE sample collection carried a pOXA-48-like plasmid, based on whether the short reads mapped to at least ≥99% of the reference plasmid obtained from the hybrid assembly of EuSCAPE_MT005. The actual proportion of isolates carrying a pOXA-48-like plasmid that belonged to one of the three high-risk lineages was calculated. The pOXA-48-like plasmids were then randomly re-distributed across all isolates in the sample collection and the proportion of pOXA-48-like plasmids in the three high-risk clonal lineages was re-calculated. This was repeated one hundred times, and the mean and 95% confidence intervals were obtained from these values.

### Replicon typing of all short-read data

Replicon typing was performed with short-read data using Ariba v2.6.1 (Hunt et al. 2017) with the PlasmidFinder database (Carattoli et al. 2014).

### Comparison of *bla*_KPC_-carrying short-read contigs with complete plasmids with NUCmer

Each short-read contig carrying a *bla*_KPC_ gene was compared with each of the complete *bla*_KPC_-carrying plasmids obtained from the hybrid assemblies using NUCmer v3.1 (Kurtz et al. 2004). Contigs that could be aligned over ≥98% of their length to a complete plasmid were deemed to match that plasmid.

### Characterisation of Tn*4401* variation

The variation within Tn*4401* and its flanking regions was characterised using TETyper v1.1 (Sheppard et al. 2018) taking the short reads of all *bla*_KPC_-carrying isolates as input.

## Data availability

All raw long-read sequence data and hybrid assemblies are available from the European Nucleotide Archive (ENA) under the study accession number, PRJEB33308 (ERP116089). Individual accession numbers for raw sequence data and hybrid assemblies are also available in **Supplementary Table 3**.

## Acknowledgements

The authors would like to thank the Pathogen Informatics team and the DNA Pipelines Long read team at the Wellcome Sanger Institute for their contribution to the study.

## Author Contributions

SD and DMA conceived the study. The EuSCAPE working group collected the bacterial isolates and epidemiological data. The ESGEM facilitated the training and capacity building for the collection of bacterial isolates. SD, VC, SR, AS, TG, JP, GMR, EJF, HG and DMA performed the data analysis. SD, GMR, EJF, HG and DMA wrote the manuscript. All authors read and approved the manuscript.

## The EuSCAPE working group

Andi Koraqi ^7^, Denada Lacej ^7^, Petra Apfalter ^8^, Rainer Hartl ^8^, Youri Glupczynski ^9^, Te-Din Huang ^9^, Tanya Strateva ^10^, Yuliya Marteva-Proevska ^11^, Arjana Tambic Andrasevic ^12^, Iva Butic ^12^, Despo Pieridou-Bagatzouni ^13^, Panagiota Maikanti-Charalampous ^13^, Jaroslav Hrabak ^14^, Helena Zemlickova ^15^, Anette Hammerum ^16^, Lotte Jakobsen ^16^, Marina Ivanova ^17^, Anastasia Pavelkovich ^17^, Jari Jalava ^18^, Monica Österblad ^18^, Laurent Dortet ^19^, Sophie Vaux ^20^, Martin Kaase ^21^, Sören G. Gatermann ^22^, Alkiviadis Vatopoulos ^23^, Kyriaki Tryfinopoulou ^23^, Ákos Tóth ^24^, Laura Jánvári ^24^, Teck Wee Boo ^25^, Elaine McGrath ^25^, Yehuda Carmeli ^26^, Amos Adler ^26^, Annalisa Pantosti ^27^, Monica Monaco ^27^, Lul Raka ^28^, Arsim Kurti ^28^, Arta Balode ^29^, Mara Saule ^29^, Jolanta Miciuleviciene ^30^, Aiste Mierauskaite ^30^, Monique Perrin-Weniger ^31^, Paul Reichert ^31^, Nina Nestorova ^32^, Sonia Debattista ^32^, Gordana Mijovic ^33^, Milena Lopicic ^33^, Ørjan Samuelsen ^34^, Bjørg Haldorsen ^34^, Dorota Żabicka ^35^, Elżbieta Literacka ^36^, Manuela Caniça ^37^, Vera Manageiro ^37^, Ana Kaftandzieva ^38^, Elena Trajkovska-Dokic ^38^, Maria Damian ^39^, Brandusa Lixandru ^39^, Zora Jelesic ^40^, Anika Trudic ^41^, Milan Niks ^42^, Eva Schreterova ^43^, Mateja Pirs ^44^, Tjasa Cerar ^44^, Jesús Oteo-Iglesias ^45^, María Pérez-Vázquez ^45^, Christian Giske ^46^, Karin Sjöström ^47^, Deniz Gür ^48^, Aslı Cakar ^48^, Neil Woodford ^49^, Katie Hopkins ^49^, Camilla Wiuff ^50^, Derek J. Brown ^51^.

^7^ University Hospital Center “Mother Theresa”, Tirana, Albania. ^8^ Elisabethinen Hospital Linz, Linz, Austria. ^9^ CHU Dinant-Godinne UCL Namur, Namur, Belgium. ^10^ Faculty of Medicine, Medical University of Sofia, Sofia, Bulgaria. ^11^ Alexandrovska University Hospital, Sofia, Bulgaria. ^12^ University Hospital for Infectious Diseases, Zagreb, Croatia. ^13^ Nicosia General Hospital, Nicosia, Cyprus. ^14^ Faculty of Medicine in Plzen, Charles University in Prague, Plzen, Czech Republic. ^15^ National Institute of Public Health, Praha, Czech Republic. ^16^ Statens Serum Institut, Copenhagen, Denmark. ^17^ East Tallinn Central Hospital, Tallinn, Estonia. ^18^ National Institute for Health and Welfare, Turku, Finland. ^19^ Bicêtre Hospital, Le Kremlin-Bicêtre, France. ^20^ Institut de Veille Sanitaire, Saint-Maurice, France. ^21^ Universitätsmedizin Göttingen, Göttingen, Germany. ^22^ Ruhr-University Bochum, Bochum, Germany. ^23^ National School of Public Health, Athens, Greece. ^24^ National Center for Epidemiology, Budapest, Hungary. ^25^ Galway University Hospitals, Galway, Ireland. ^26^ Ministry of Health, Tel-Aviv, Israel. ^27^ Istituto Superiore di Sanità, Rome, Italy. ^28^ National Institute of Public Health of Kosovo, Prishtina, Kosovo. ^29^ Pauls Stradins Clinical University Hospital, Riga, Latvia. ^30^ National Public Health Surveillance Laboratory, Vilnius, Lithuania. ^31^ Laboratoire National De Sante, Düdelingen, Luxembourg. ^32^ Mater Dei Hospital, Msida, Malta. ^33^ Institute of Public Health, Podgorica, Montenegro. ^34^ University Hospital of North Norway, Tromsø, Norway. ^35^ Narodowy Instytut Lekow, Warsaw, Poland. ^36^ National Medicines Institute, Warsaw, Poland. ^37^ National Institute of Health Dr. Ricardo Jorge, Lisbon, Portugal. ^38^ Institute of Microbiology and Parasitology, Medical Faculty, Skopje, Republic of Macedonia. ^39^ Cantacuzino National Research Institute, Bucharest, Romania. ^40^ Institute for Public Health of Vojvodina, Novi Sad, Serbia. ^41^ Institute for Pulmonary Diseases of Vojvodina, Sremska Kamenica, Serbia. ^42^ Public Health Authority of the Slovak Republic, Bratislava, Slovakia. ^43^ University Hospital of P.J.Safarik, Kosice, Slovakia. ^44^ Institute of Microbiology and Immunology, Ljubljana, Slovenia. ^45^ Centro Nacional de Microbiología, Instituto de Salud Carlos III, Madrid, Spain. ^46^ Karolinska Institutet, Stockholm, Sweden. ^47^ Public Health Agency of Sweden, Stockholm, Sweden. ^48^ Hacettepe University, Ankara, Turkey. ^49^ National Infection Service, Public Health England, London, United Kingdom - England and Northern Ireland. ^50^ Sydvestjysk Hospital, Esbjerg, Denmark. ^51^ Scottish Microbiology Reference Laboratories, Glasgow, United Kingdom - Scotland

