## Supplementary Information for "Genomic analysis of carbapenemase-encoding plasmids from *Klebsiella pneumoniae* across Europe highlights three major patterns of dissemination"

**Supplementary Figures**

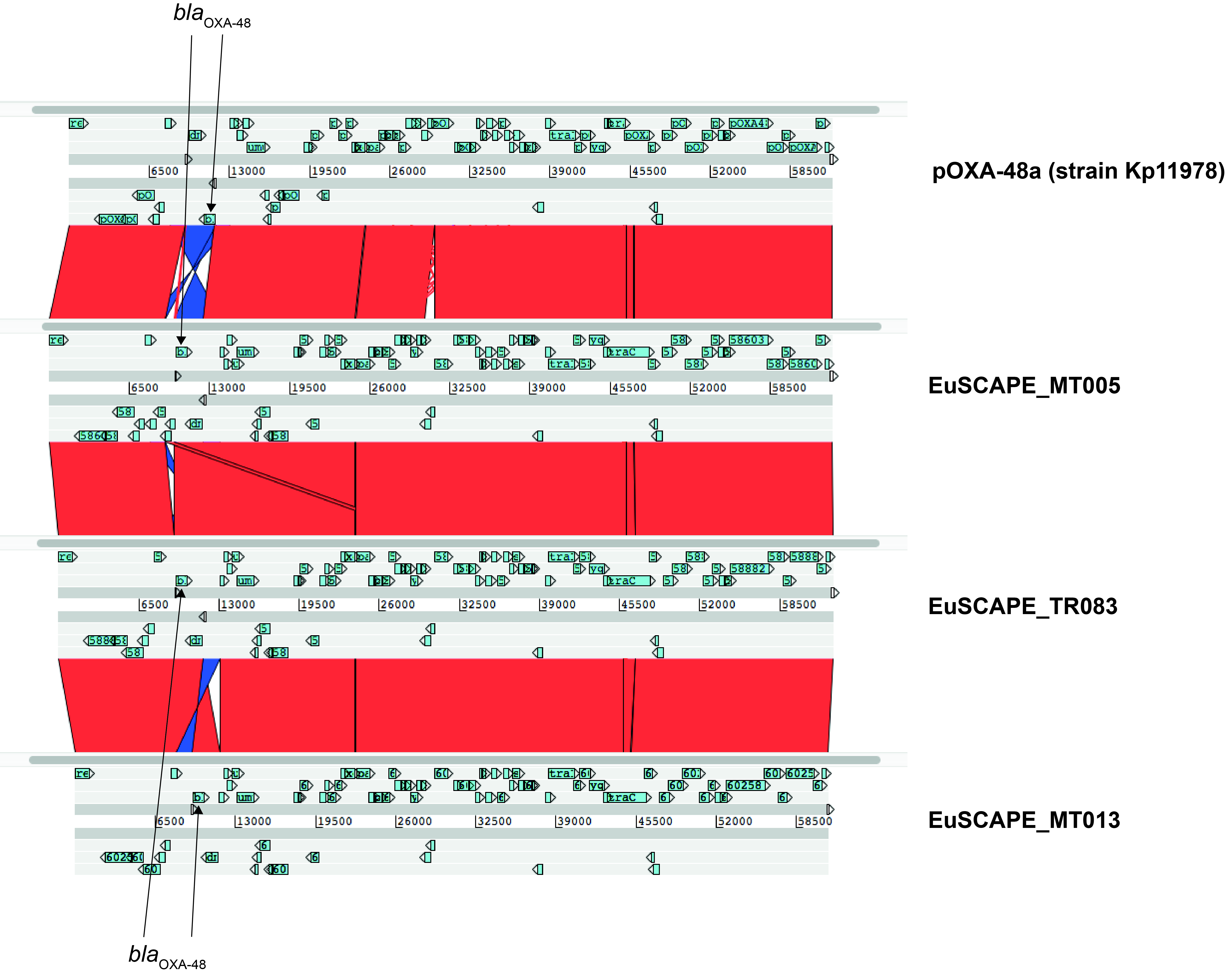

**Supplementary Figure 1. High structural similarity between *bla*_OXA-48-like_-carrying pOXA-48-like plasmids.** A comparison with the Artemis Comparison Tool (ACT) shows the high structural homology between pOXA-48a (Poirel et al. 2012) and plasmids from the hybrid assemblies of EuSCAPE_MT005 (circularised), EuSCAPE_TR083 (circularised) and EuSCAPE_MT013 (non-circularised). Sections of similarity of ≥500bp are shown.

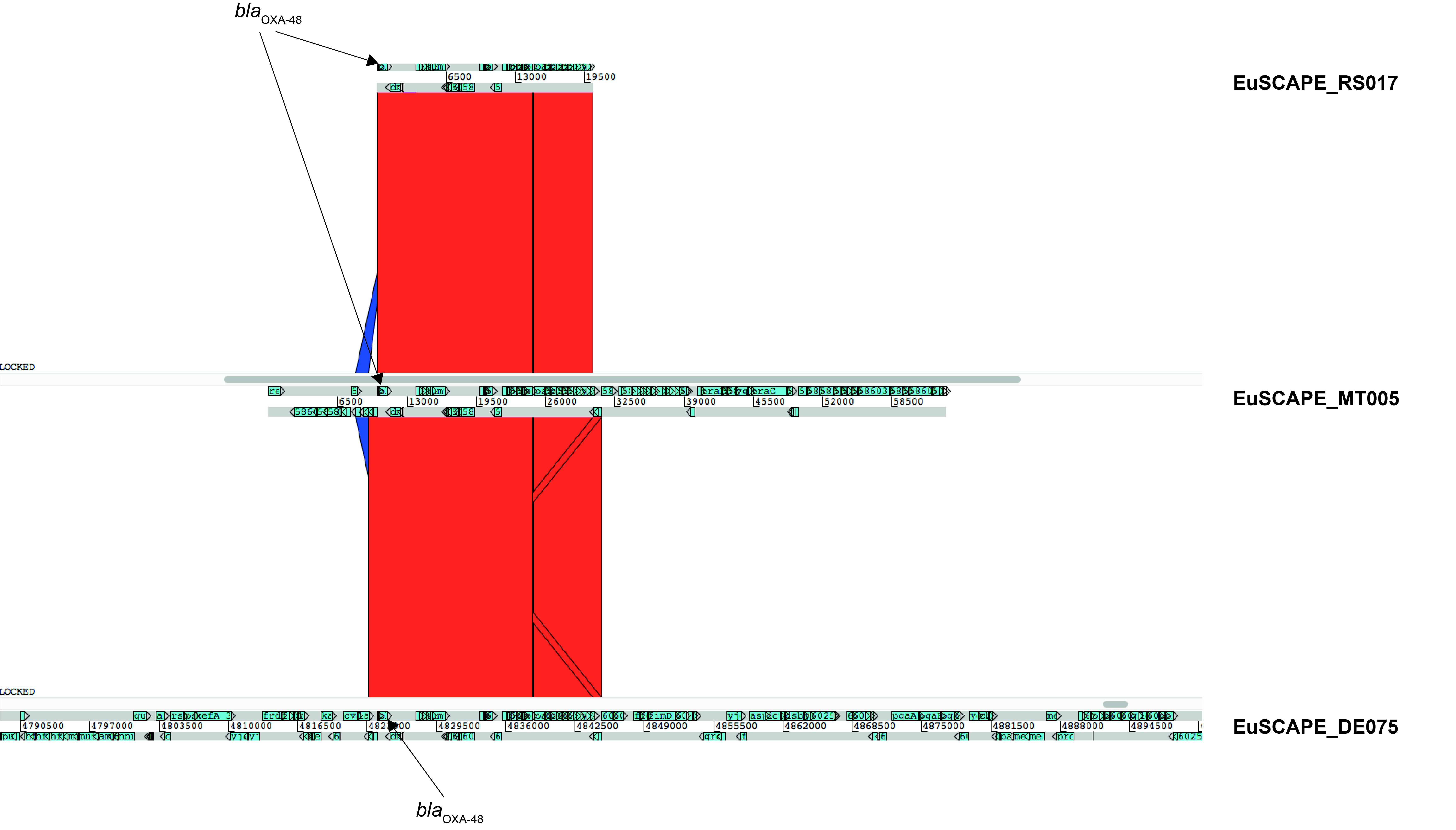

**Supplementary Figure 2. Integration of the Tn*6237* transposon carrying *bla*_OXA-48_ into the chromosome.** A comparison with the Artemis Comparison Tool (ACT) shows the homology between different vectors of *bla*_OXA-48_. From top to bottom are the 20.3kb non-circularised putative plasmid sequence from EuSCAPE_RS017 which represents the Tn*6237* transposon, the 63.5kb circularised putative plasmid sequence from EuSCAPE_MT005 which represents a full-length pOXA-48-like plasmid, and a section of the chromosomal sequence of EuSCAPE_DE075 into which the *bla*_OXA-48_ gene has integrated. Sections of similarity of ≥500bp are shown.

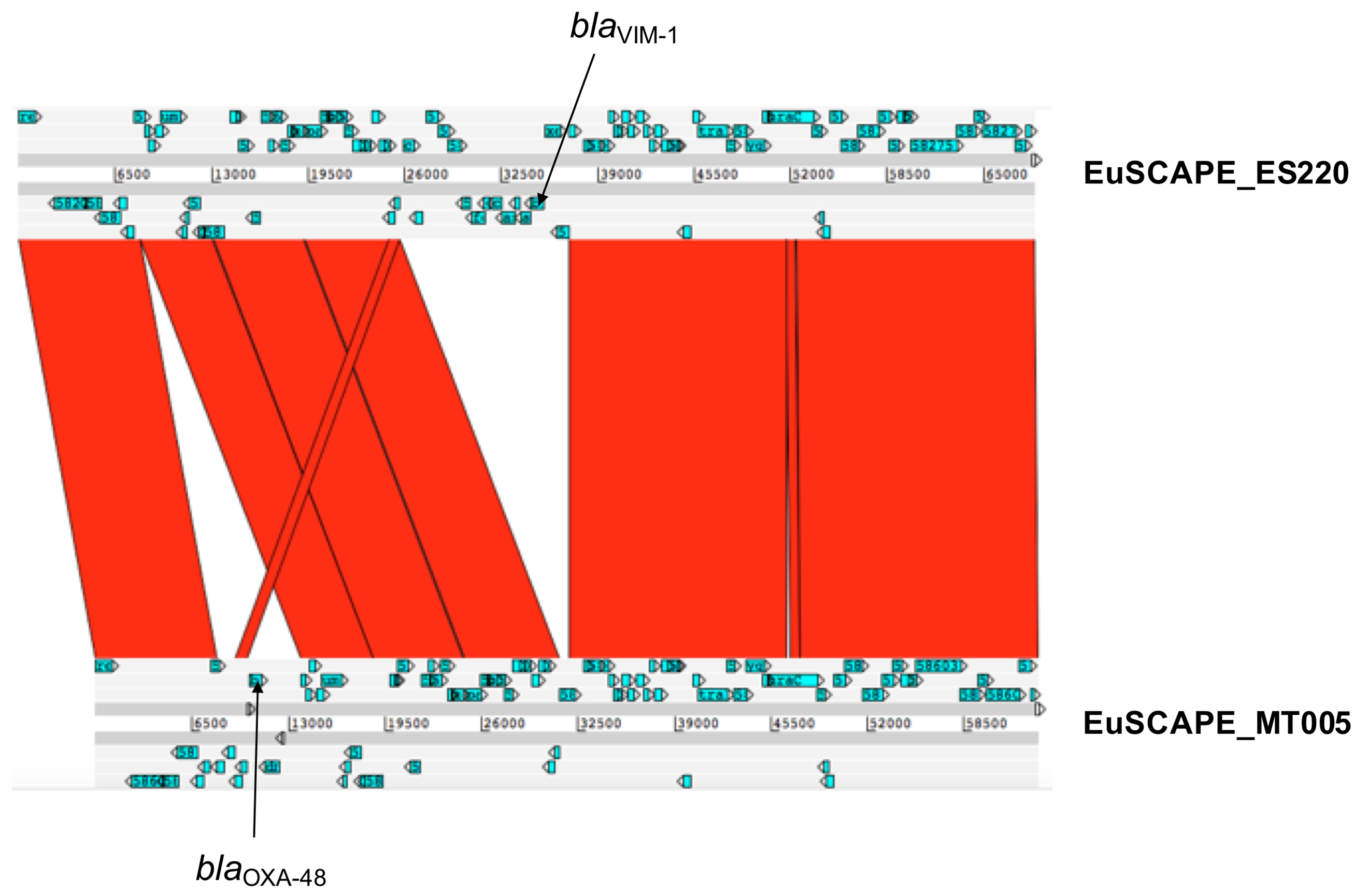

**Supplementary Figure 3. Integration of *bla*_VIM-1_ into the pOXA-48-like plasmid.** A comparison with the Artemis Comparison Tool (ACT) between the *bla*_VIM-1_-carrying plasmid from EuSCAPE_ES220 and the *bla*_OXA-48_-carrying (pOXA-48-like) plasmid from EuSCAPE_MT005 shows high structural homology, with differences only in the presence/absence of mobile regions carrying *bla*_OXA-48_ and *bla*_VIM-1_. Both plasmids could be circularised. Sections of similarity of ≥500bp are shown.

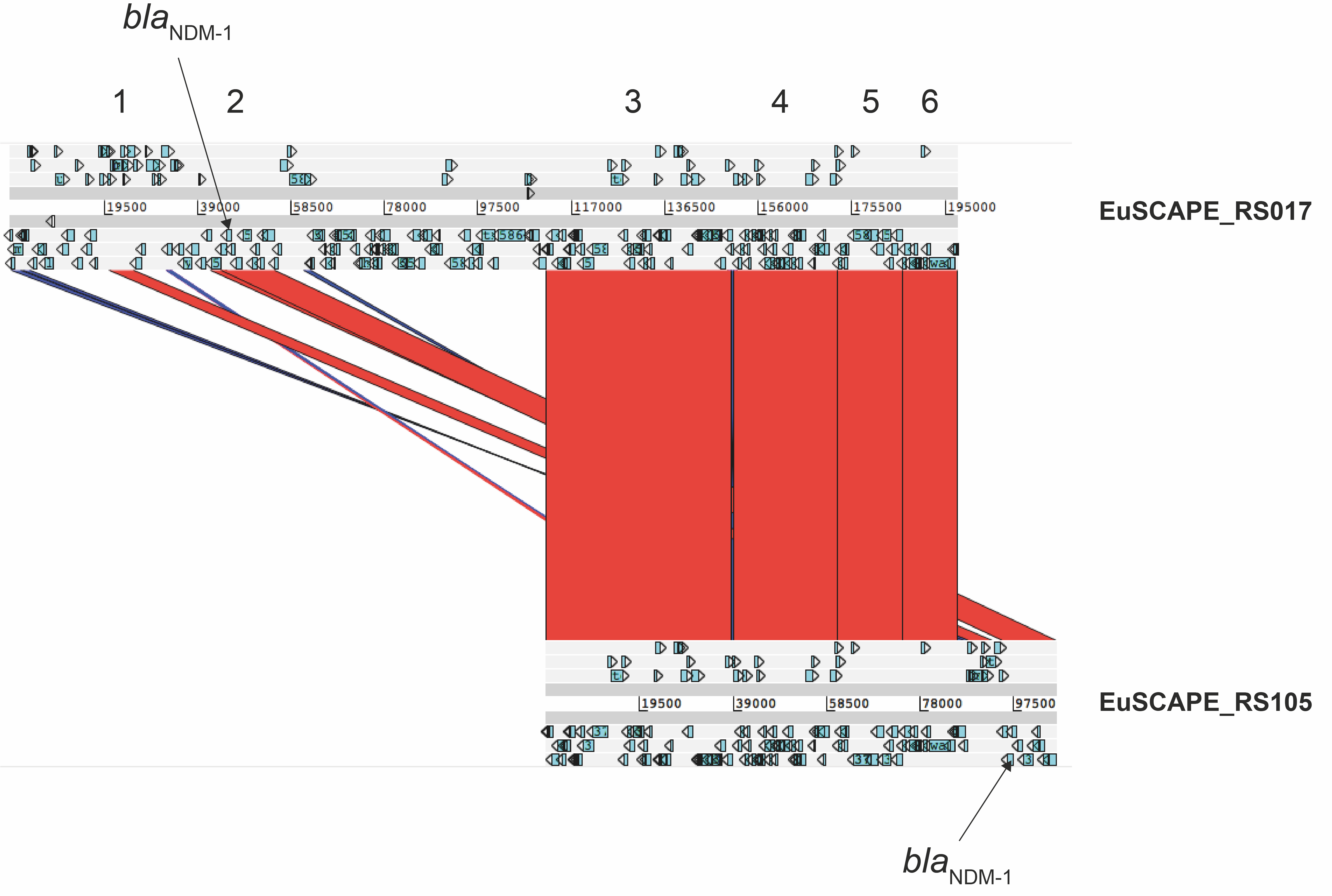

**Supplementary Figure 4. Recent shared ancestry between *bla*_NDM-1_-carrying plasmids from ST274 and ST101.** A comparison is shown between the *bla*_NDM-1_-carrying plasmids from EuSCAPE_RS105 (ST274) and EuSCAPE_RS017 (ST101) that harbour an IncA/C2 replicon and both IncA/C2 and IncR replicons, respectively. Neither plasmid could be circularised. The nucleotide similarity across six major shared regions is high, as demonstrated by the number of bases and SNPs found in each: 1) 4,958bp, 1 SNP; 2) 11,180bp, 1 SNP; 3) 38,626bp, 1 SNP; 4) 21,560, 0 SNPs; 5) 13,600bp, 0 SNPs; 6) 11,500bp, 0 SNPs.

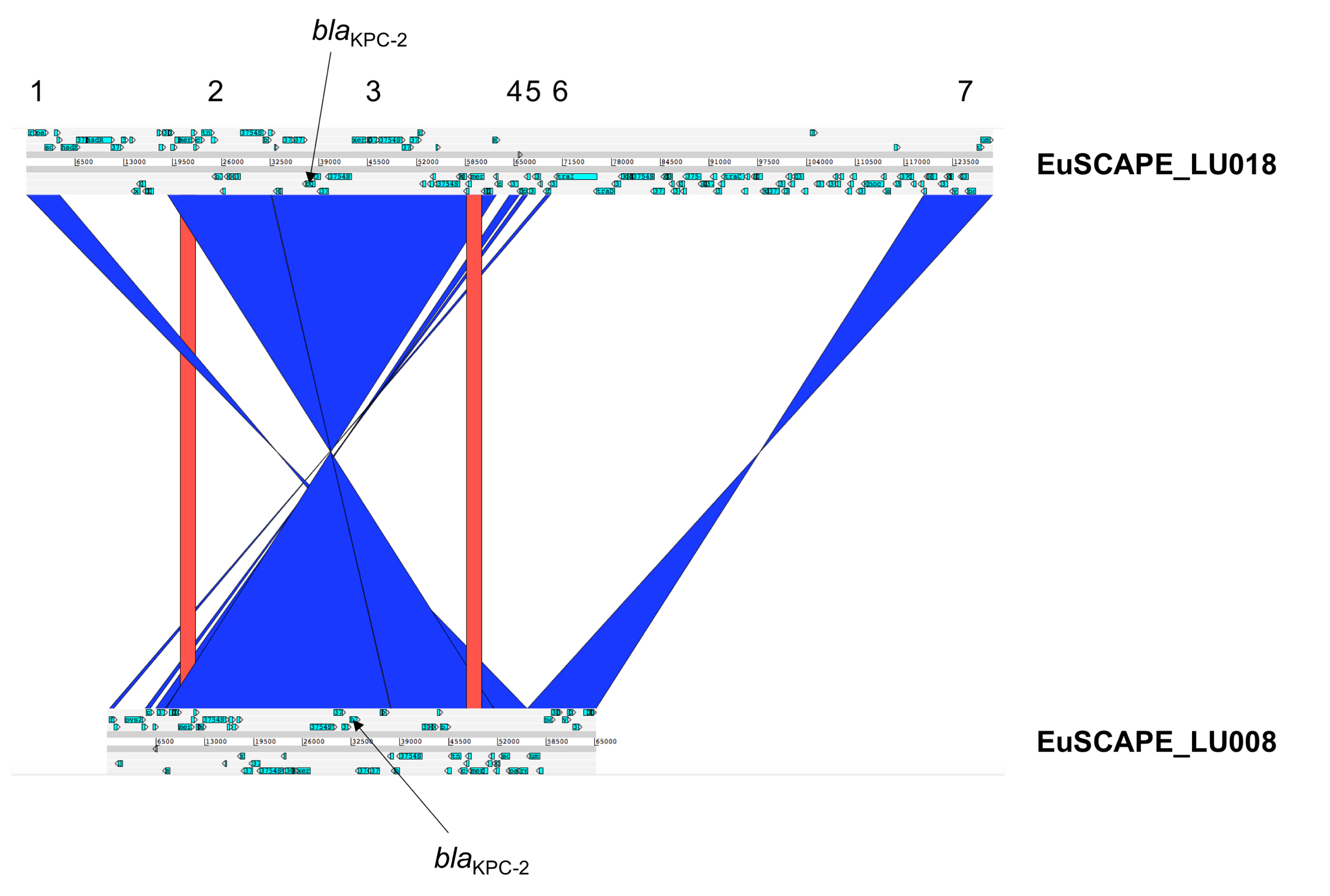

**Supplementary Figure 5. Evidence for recent common ancestry between two plasmids obtained from isolates of different STs but from the same hospital.** A comparison with the Artemis Comparison Tool (ACT) between the *bla*_KPC-2_-carrying plasmid sequences from EuSCAPE_LU018 (circularised) and EuSCAPE_LU008 (non-circularised) shows seven regions of shared sequence (≥500bp). The nucleotide similarity across these regions differs substantially (suggestive of different divergence times), as demonstrated by the number of bases and SNPs found in each: 1) 4,374bp, 0 SNPs; 2) 13,741bp, 0 SNPs; 3) 30,013bp, 0 SNPs; 4) 1,286bp, 25 SNPs; 5) 693bp, 44 SNPs; 6) 563bp, 26 SNPs; 7) 9,174bp, 0 SNPs.

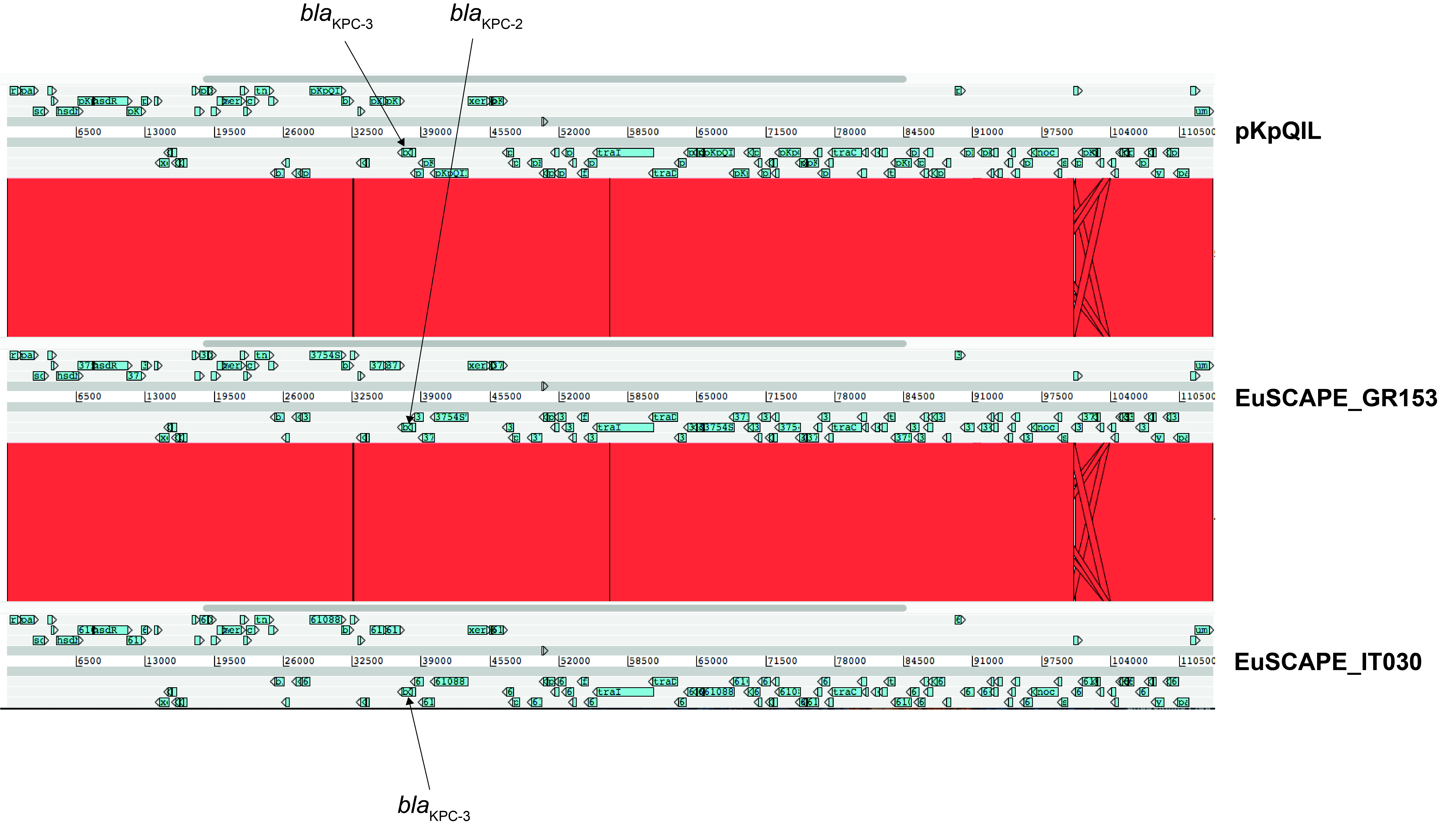

**Supplementary Figure 6. High structural similarity between *bla*_KPC_-carrying pKpQIL-like (backbone I) plasmids.** A comparison with the Artemis Comparison Tool (ACT) shows the high structural homology between pKpQIL (Leavitt et al. 2010) and circularised plasmids from EuSCAPE_GR153 and EuSCAPE_IT030. Sections of similarity of ≥500bp are shown.

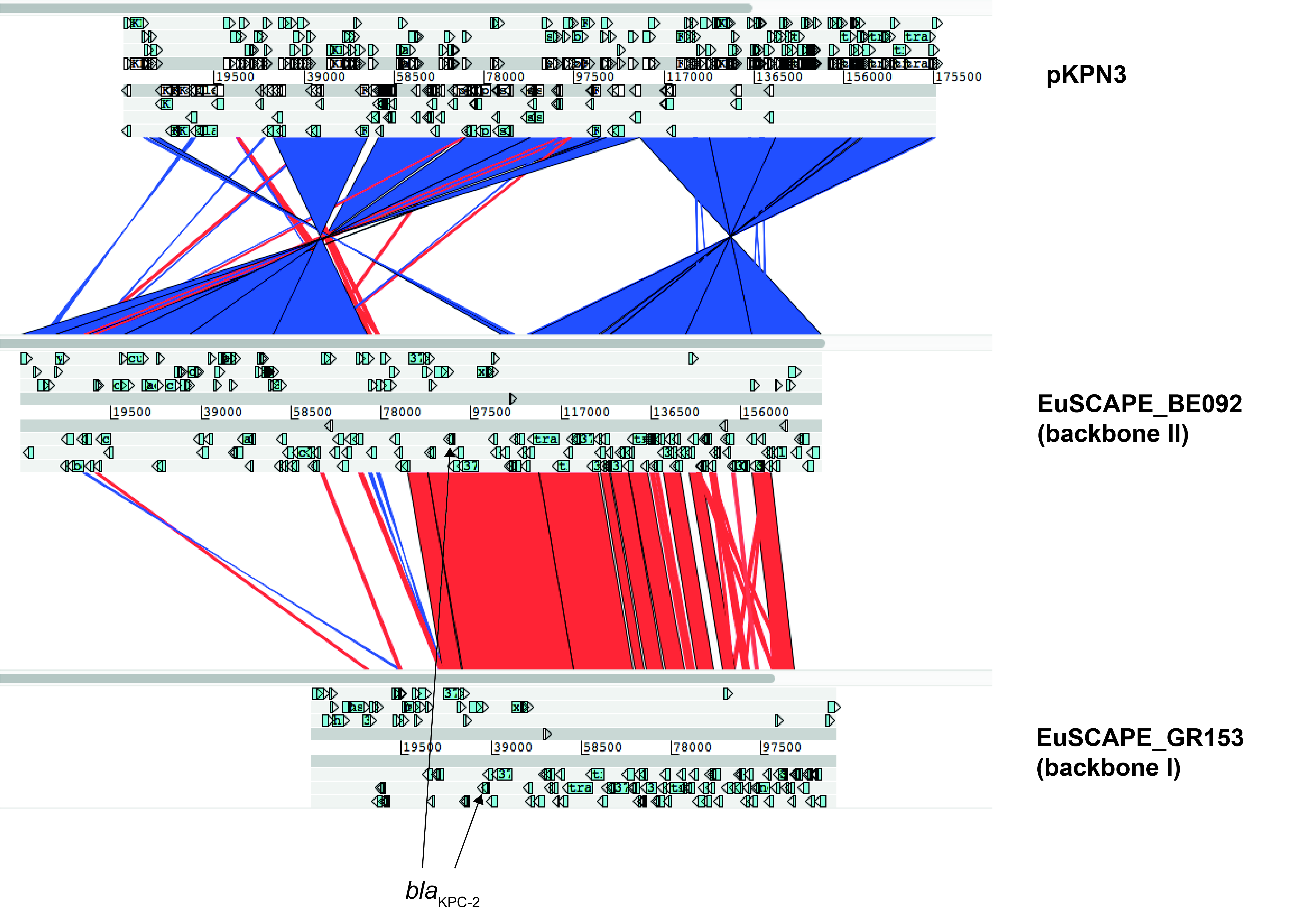

**Supplementary Figure 7. Partial sequence homology between pKPN3 and IncF plasmid backbones I and II that carry *bla*_KPC_.** A comparison with the Artemis Comparison Tool (ACT) between pKPN3 (accession number, CP000648) and the circularised *bla*_KPC-2_-carrying plasmids of EuSCAPE_GR153 (backbone I) and EuSCAPE_BE092 (backbone II) is shown. Sections of similarity of ≥500bp are shown.

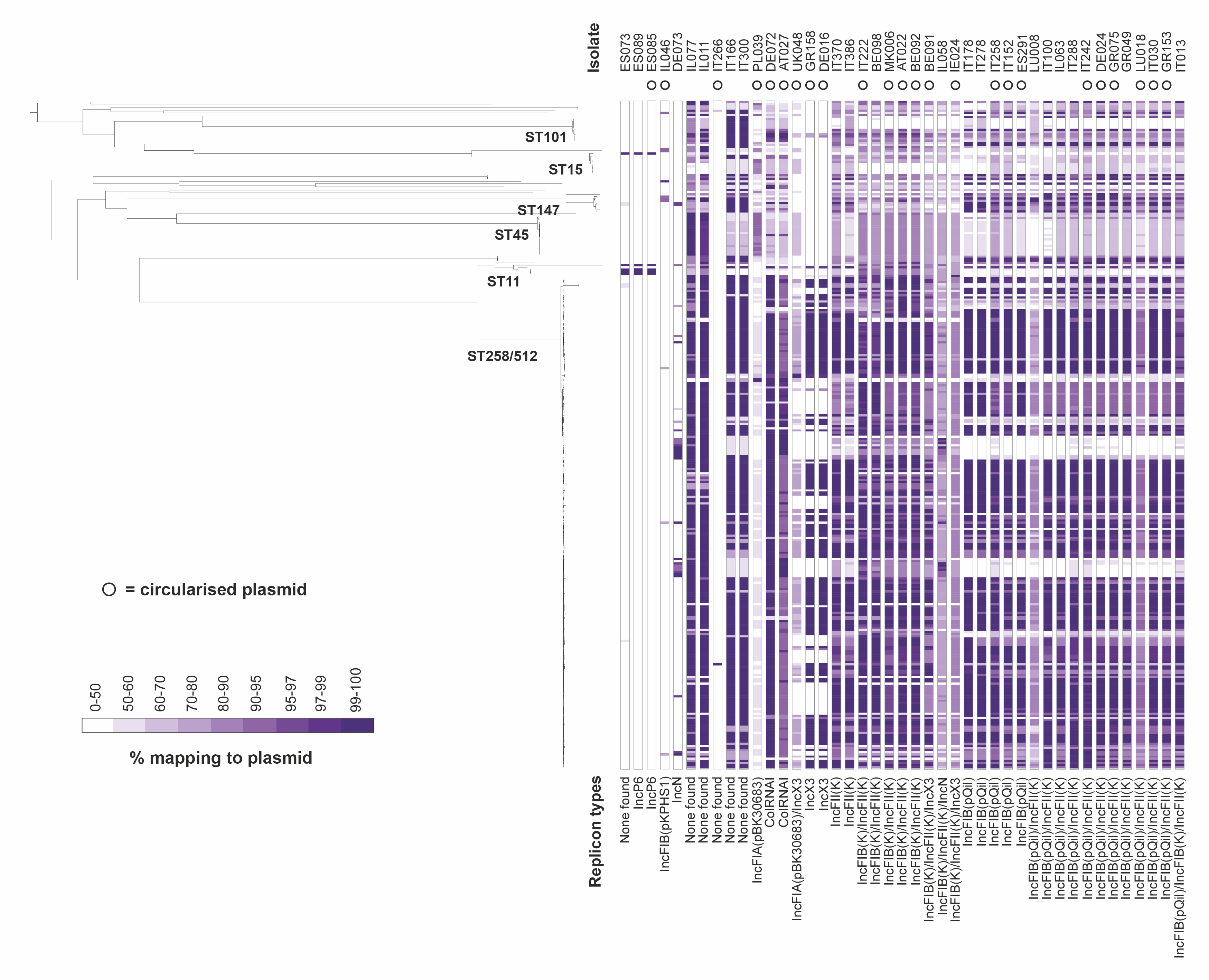

**Supplementary Figure 8. Isolates carrying *bla*_KPC_ genes frequently carry two or more of the different plasmid types found to harbour *bla*_KPC_ genes in the hybrid assemblies.** The phylogenetic tree shows 311 *bla*_KPC_-carrying *K. pneumoniae sensu stricto* isolates (the *bla*_KPC_-carrying isolate from *K. variicola* is excluded). The tree was constructed using SNPs in the core genome and is midpoint rooted. All non-*bla*_KPC_-carrying isolates, which would be interspersed amongst the isolates here, were also excluded. The columns show the percentage lengths of each of the 43 *bla*_KPC_-carrying putative plasmid sequences found in the hybrid assemblies that were mapped by the short reads of the 311 *bla*_KPC_-carrying isolates (note the non-linear colour gradient). The replicon types found in each of the plasmid sequences are indicated at the bottom.

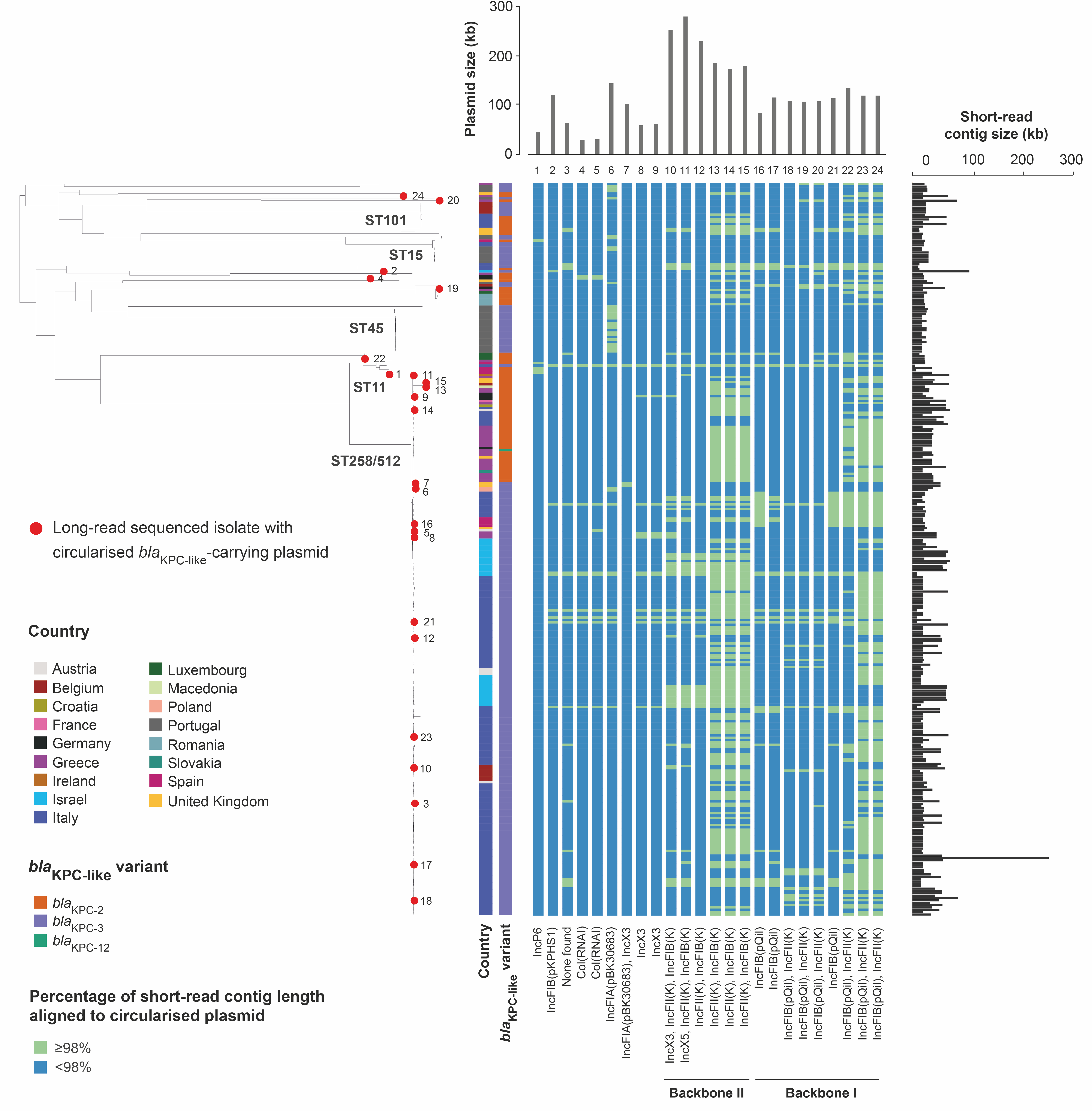

**Supplementary Figure 9. Most *bla*_KPC_-carrying contigs from the short-read assemblies are compatible with belonging to IncF backbone I and II plasmids only.** The phylogenetic tree contains 311 *bla*_KPC_-carrying *K. pneumoniae sensu stricto* isolates (the *bla*_KPC_-carrying isolate from *K. variicola* is excluded). The tree was constructed using SNPs in the core genome and is midpoint rooted. All non-*bla*_KPC_-carrying isolates, which would be interspersed amongst the isolates here, were also excluded. Twenty-four isolates from which circularised *bla*_KPC_-carrying plasmids were obtained are marked by red circles in the tree and numbered. The columns, from left to right, show the country of isolation, the *bla*_KPC_ variant, and whether the short-read assembly contigs carrying *bla*_KPC_ are compatible with each of the 24 complete plasmids (i.e. ≥98% of the contig aligns to the plasmid using NUCmer). The latter columns are numbered according to the isolate from which each plasmid was recovered from in the phylogenetic tree (and in the same order as in the matrix of Figure 3). The length of the circularised plasmids and the short-read assembly contigs are shown.

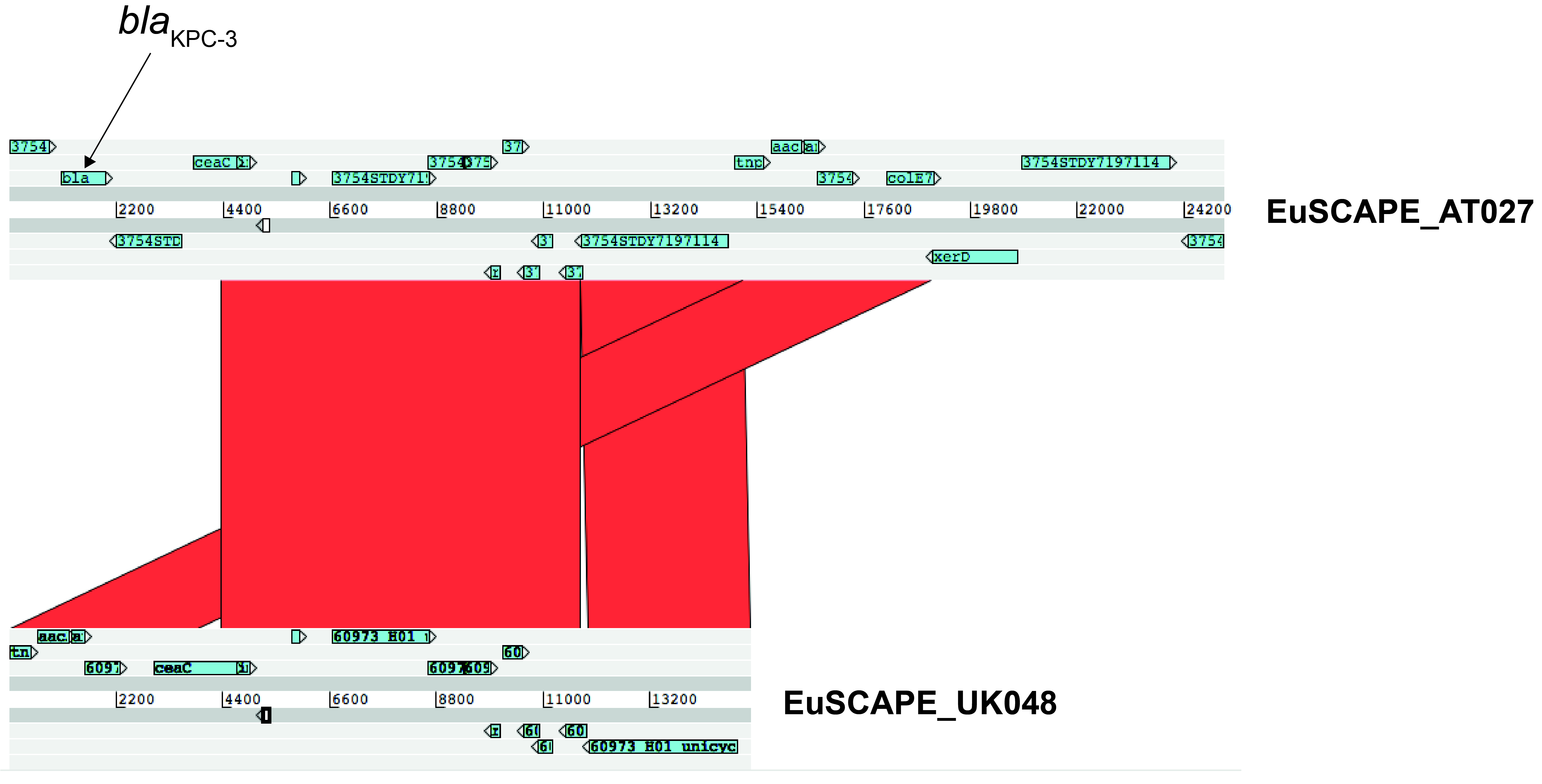

**Supplementary Figure 10. Structural homology of ColRNAI plasmids with and without *bla*_KPC_ genes.** The structural similarity between the *bla*_KPC-3_-carrying plasmid sequence from EuSCAPE_AT027 and non-*bla*_KPC_-carrying plasmid sequence from EuSCAPE_UK048 is visualized using the Artemis Comparison Tool (ACT). Both plasmids could be circularised. Sections of similarity of ≥500bp are shown.

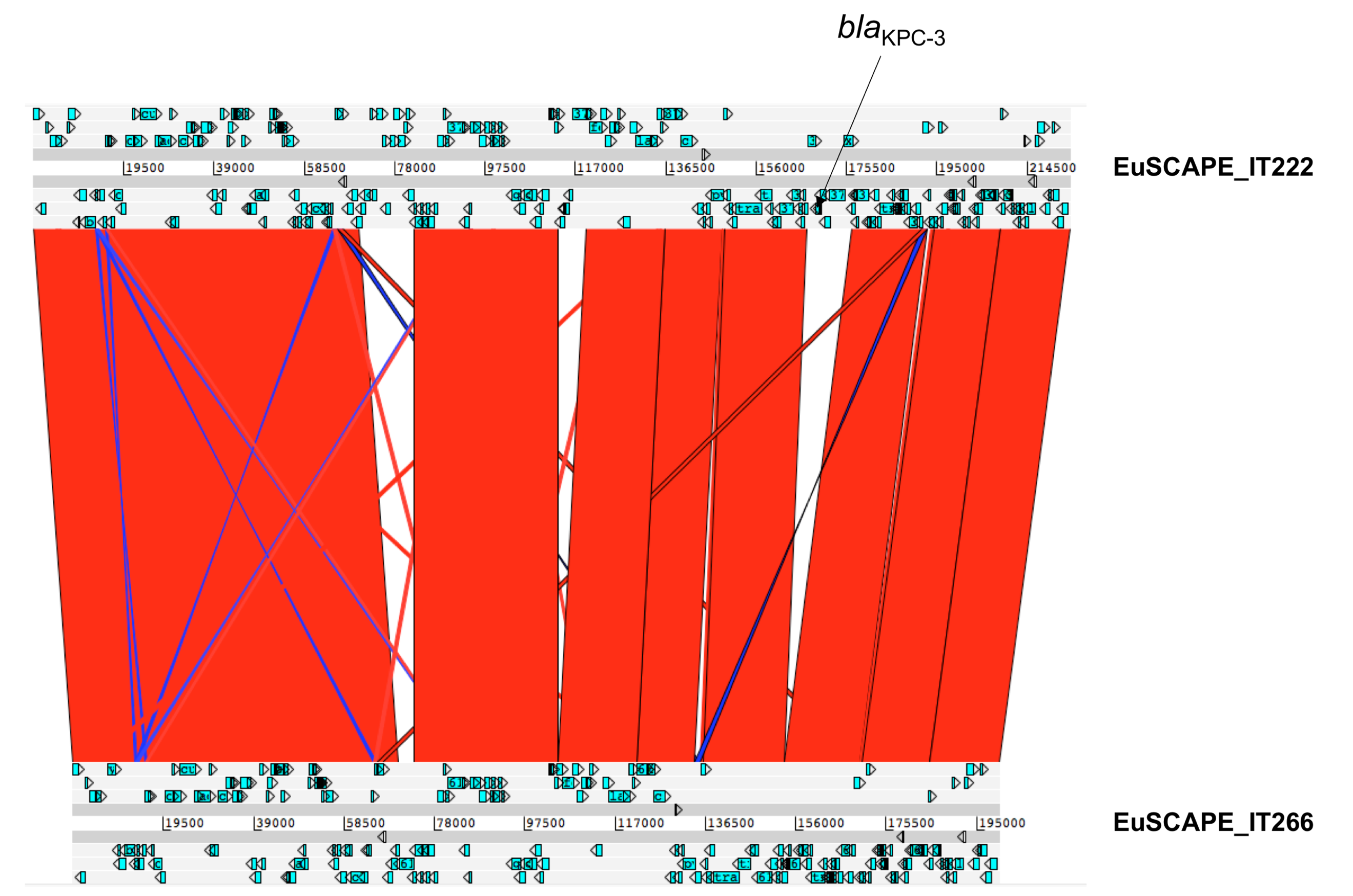

**Supplementary Figure 11. Structural homology of backbone II (IncFIB(K)/IncFII(K)) plasmids with and without *bla*_KPC_ genes.** The structural similarity between the *bla*_KPC-3_-carrying plasmid sequence from EuSCAPE_IT222 and non-*bla*_KPC_-carrying plasmid sequence from EuSCAPE_IT266 is visualized using the Artemis Comparison Tool (ACT). Both plasmids could be circularised. Sections of similarity of ≥500bp are shown.

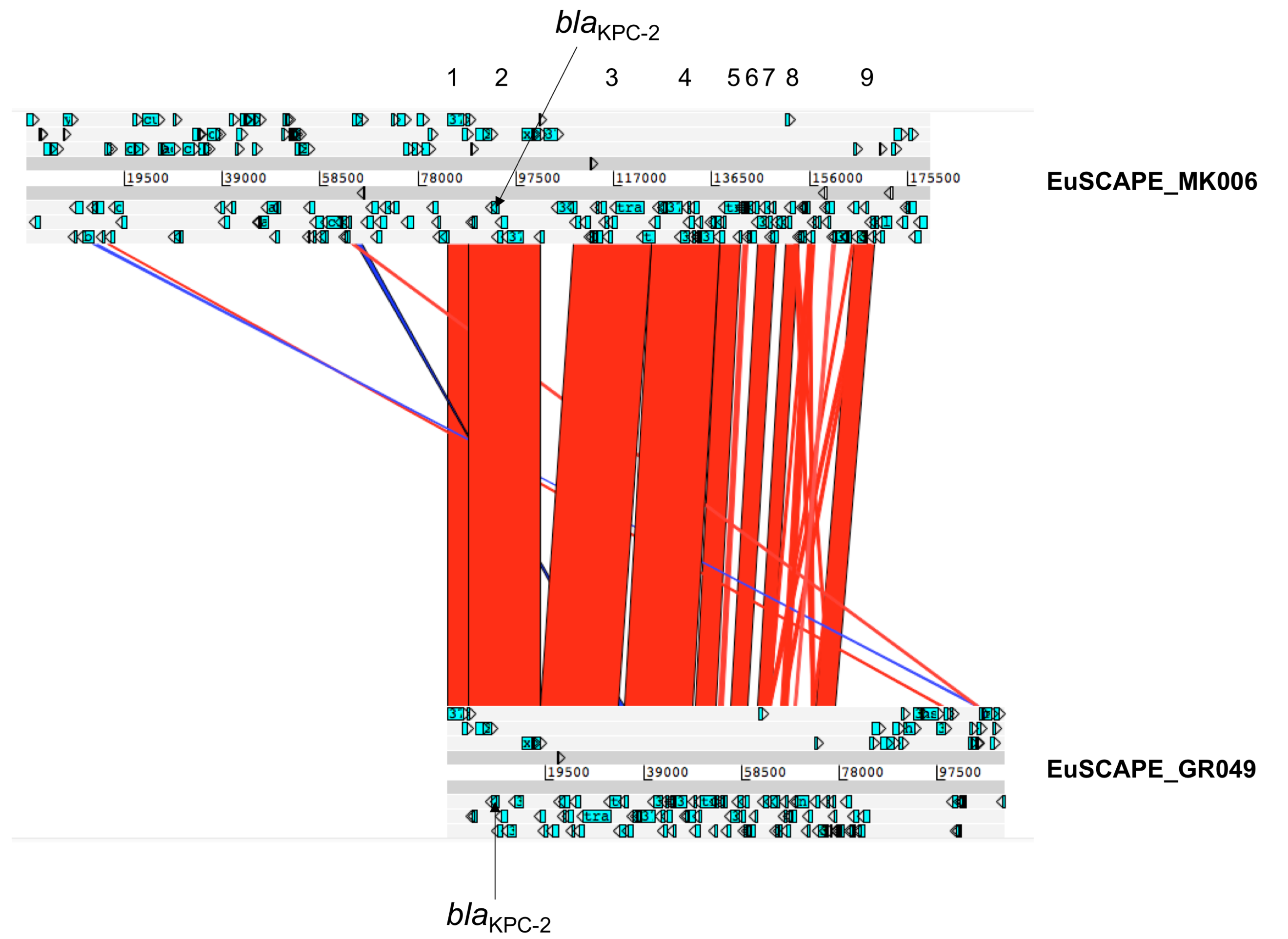

**Supplementary Figure 12. High nucleotide similarity over 34kb between two plasmids representing backbones I and II suggests recombination involving the *bla*_KPC_ gene.** The structural similarity between the *bla*_KPC-2_-carrying plasmid sequences from EuSCAPE_GR049 (non-circularised) and EuSCAPE_MK006 (circularised) is visualized using the Artemis Comparison Tool (ACT). These are from closely-related isolates and represent backbones I and II of the *bla*_KPC_-carrying plasmids, respectively. Sections of similarity of ≥500bp are shown. High nucleotide similarity was found in three of nine highlighted shared regions which comprise a total of 34kb. The total number of bases and SNPs found in the nine regions are: 1) 4059bp, 0 SNPs; 2) 14,293bp, 2 SNPs; 3) 15,671bp, 0 SNPs; 4) 13,574bp, 53 SNPs; 5) 4150bp, 109 SNPs; 6) 1012bp, 75 SNPs; 7) 3600bp, 91 SNPs; 8) 2606bp, 58 SNPs; 9) 3986bp, 61 SNPs.

**Supplementary Note**

**Inferring a chromosomal or plasmid origin of contigs in the hybrid assemblies**

11/79 (13.9%) putative plasmid sequences carrying carbapenemase genes in the hybrid assemblies could be neither circularised nor contained plasmid replicons. Visualisation of the assembly graphs using Bandage (Wick et al. 2015) suggested that 10/11 belong to plasmids since they connect with other non-chromosomal contigs. Furthermore, we found 4/11 were from hybrid assemblies with a circularised chromosomal sequence, which also lends support to a plasmid origin of the remaining smaller sequences. Finally, we investigated whether we could use sequencing coverage of the carbapenemase genes from the short-read data to infer their origin. We found that the short-read coverage of 41 carbapenemase genes found on circularised contig sequences in the hybrid assemblies with known plasmid replicons (i.e. sequences that we were highly confident represented plasmids) was highly variable relative to multi-locus sequence typing (MLST) genes (0.3x–27.8x) and had a median value of 1.7x. The median coverage (relative to MLST genes) of the five carbapenemase genes found on chromosomal sequences was identical to this, although the values were less variable (1.4-1.9x). These results suggest that high coverage of a carbapenemase gene may indicate a plasmid origin but low coverage could represent either a plasmid or chromosomal origin. Based on this, we assumed a plasmid origin for 4/11 of the putative plasmid sequences that had ≥3x coverage of the carbapenemase (relative to MLST genes) in the short-read data. Only one of these overlapped with the four associated with a circularised chromosome.

**Comparison of IncL/M(pOXA48) plasmids with pOXA-48a**

Using NUCmer (Kurtz et al. 2004), we found 0-4 SNPs between three IncL/M(pOXA48) plasmids carrying *bla*_OXA-48_ that were obtained from hybrid assemblies (EuSCAPE_MT013, EuSCAPE_MT005, EuSCAPE_TR083). We found 128 SNPs between each of these three plasmids and the previously published plasmid, pOXA-48a (Poirel et al. 2012). Most of these SNPs were highly clustered, suggesting that they have been introduced by recombination. The four plasmids showed high structural homology using the Artemis Comparison Tool (ACT) (Carver et al. 2005) (**Supplementary Figure 1**).

**Most *bla*_OXA-48-like_-carrying plasmids were found rarely**

Using short-read mapping to *bla*_OXA-48-like_-carrying plasmids obtained from the hybrid assemblies, we found that the ColKP3, IncX3, IncL/M(pMU407) and IncA/C2 plasmid sequences were found only rarely amongst the 249 *bla*_OXA-48-like_-carrying isolates (**Figure 1A**). In total, 6 (2.4%), 3 (1.2%), 3 (1.2%) and 1 (0.4%) isolate(s), including the long-read sequenced isolates themselves, had mapping across ≥99% of each of the four plasmid types, respectively. These numbers only increased to 6 (2.4%), 3 (1.2%), 12 (4.8%) and 16 (6.4%) when the mapping threshold was lowered from ≥99% to ≥90%.

**Chromosomal integration of *bla*_OXA-48_ genes**

We found *bla*_OXA-48_ within the chromosome of three hybrid assemblies recovered from two closely-related ST11 isolates (EuSCAPE_ES046 and EuSCAPE_ES089) and one ST530 isolate (EuSCAPE_DE075). Within all three chromosomes, *bla*_OXA-48_ was found within a ~20kb composite transposon, Tn*6237*, which also carried *bla*_OXA-48_ in pOXA-48-like plasmids (**Supplementary Figure 2**). The Tn*6237* transposon (carrying *bla*_OXA-48-like_) was found in the same chromosomal location in both ST11 assemblies, indicative of a single integration event. We also found direct repeat sequences (CAACCGGCA – EuSCAPE_ES046/EuSCAPE_ES089; CGCTCGAG – EuSCAPE_DE075) immediately upstream and downstream of each transposon sequence, which likely facilitate integration.

To further confirm the chromosomal integration of *bla*_OXA-48_ in these three isolates, we mapped their long reads to the chromosome from each of their own hybrid assemblies. We found reads bridging the junction between the chromosomal sequence and integrated Tn*6237* transposon, supporting chromosomal integration. By contrast, when the long reads of isolates with full-length pOXA-48-like plasmids were mapped to the chromosomes of EuSCAPE_ES046, EuSCAPE_ES089 and EuSCAPE_DE075, we found an absence of reads spanning the chromosome/transposon junction.

Using the short-read sequence data, we also found evidence of other closely-related isolates carrying *bla*_OXA-48_ genes within the chromosome. Two of the three isolates with a chromosomally integrated *bla*_OXA-48_ gene, EuSCAPE_ES046 and EuSCAPE_ES089, belonged to a clade of eight closely-related *bla*_OXA-48_-carrying ST11 isolates submitted from three different hospitals in Madrid, Spain. The short sequence reads of all eight of these isolates mapped to 100% of the Tn*6237* transposon. However, they mapped to various extents (36.3-100%) of the full-length 63.5kb pOXA-48-like plasmid from EuSCAPE_MT005 (i.e. with at least one read per base) (**Figure 1A**). In particular, short reads from EuSCAPE_ES046 and EuSCAPE_ES089 mapped to 47.3% and 94.5% of the 63.5kb epidemic plasmid, respectively, suggesting that they had retained additional pOXA-48-like plasmid sequence despite integration of the Tn*6237* transposon into the chromosome. However, we inspected the mapped sequence reads of all eight isolates to the 63.5kb plasmid using Artemis (Carver et al. 2012), and found that 7/8 isolates had high (>50x) and even coverage across the ~20kb Tn*6237* region, but either no or very low coverage of reads (typically <5x) across the remaining sequence. This supports chromosomal integration of *bla*_OXA-48_ in these isolates. However, one isolate, EuSCAPE_ES068, had even coverage (~200x) across the entire pOXA-48-like plasmid, suggesting it still possessed an independent full-length plasmid.

The third hybrid assembly with a chromosomally-integrated *bla*_OXA-48_ gene is from EuSCAPE_DE075, which belonged to a clade of three closely-related ST530 isolates. We found that, in addition to EuSCAPE_DE075, another of these also may have carried *bla*_OXA-48_ within the chromosome. Short-read mapping showed that it possessed the full ~20kb Tn*6237* sequence, but only 46.0% of the 63.5kb pOXA-48-like plasmid sequence.

Finally, we also found that a 20.3kb non-circularised putative plasmid sequence carrying *bla*_OXA-48_ from the hybrid assembly of EuSCAPE_RS017 (ST101) represented the Tn*6237* transposon. Another 24.9kb of sequence from the pOXA-48-like plasmid which contains the IncL/M(pOXA48) replicon was found in the same assembly on a different contig. Thus the pOXA-48-like plasmid may have been present in this isolate although there could also have been recent movement of Tn*6237* out of the plasmid, resulting in the separate 20.3kb fragment.

**Integration of *bla*_VIM-1_ into a pOXA-48-like plasmid**

Amongst the hybrid assemblies, we found a 68.4kb IncL/M(pOXA48) plasmid carrying *bla*_VIM-1_. This was recovered from a ST483 isolate, EuSCAPE_ES220. The plasmid had an identical backbone structure to pOXA-48-like plasmids, except for a deletion of the 4.9kb Tn*1999* region containing *bla*_OXA-48_ and insertion of a 12.2kb Tn*21* region containing *bla*_VIM-1_ at a different location (**Supplementary Figure 3**). Short sequence reads from a further four *bla*_VIM-1_-carrying isolates, none of which carried a *bla*_OXA-48-like_ gene, also mapped to the full length of the 68.4kb *bla*_VIM-1_-carrying plasmid. These comprised another ST483 isolate that was very closely related to EuSCAPE_ES220, as well as one ST11 and two ST15 isolates.

We next determined the evolutionary relationships between pOXA48-like plasmids carrying *bla*_OXA-48_ and *bla*_VIM-1_ genes. We mapped the short sequence reads of EuSCAPE_ES220, together with those from the other four *bla*_VIM-1_-carrying isolates, to the *bla*_OXA-48_-carrying plasmid from EuSCAPE_MT005. Combined phylogenetic analyses of these with all other pOXA-48-like plasmids carrying *bla*_OXA-48-like_ genes showed that four of the five (putative) *bla*_VIM-like_-carrying plasmids cluster monophyletically (including EuSCAPE_ES220) (**Figure 1B**). These four mapped plasmid sequences have pairwise SNP differences of 0-1. Together, these results further support the presence of *bla*_VIM-1_ on the pOXA-48-like plasmid backbone for three of the non-long-read sequenced isolates (1 x ST483, 2 x ST15). It also suggests that integration of the 12.2kb region carrying *bla*_VIM-1_ occurred once in a common ancestor of the four monophyletic plasmid sequences, and that the plasmid has since spread horizontally at least once (i.e. between ST15 and ST483). The fifth putative *bla*_VIM-1_-carrying plasmid sequence from an ST11 isolate differs from these by 58-59 SNPs, and is also more related to other pOXA-48-like plasmid sequences harbouring *bla*_OXA-48-like_ genes. This suggests that if *bla*_VIM-1_ is on a pOXA-48-like backbone in this isolate, the gene integrated on a separate occasion.

**Comparison of *bla*_KPC_-carrying plasmids from isolates belonging to different STs but submitted from the same hospital**

We generated hybrid assemblies for two isolates (EuSCAPE_LU018, EuSCAPE_LU008) belonging to the same *bla*_KPC_ genetic context (GC) group. This was to investigate possible plasmid transfer since they were submitted from the same hospital but belonged to different STs. We obtained a 128.8kb circularised plasmid from EuSCAPE_LU018 and a non-circularised 65.1kb plasmid from EuSCAPE_LU008, which carried *bla*_KPC_ genes. Both were backbone I plasmids harbouring IncFII(K) and IncFIB(pQIL) replicons. A comparison with the Artemis Comparison Tool (Carver et al. 2005) showed structural rearrangements between the two (**Supplementary Figure 5**). Despite this, we found 0 SNPs across 95.8% of the total shared sequence. This is suggestive of a recent common ancestor and therefore of possible plasmid exchange within the hospital.

**Comparison of short-read contigs from ST258/512 isolates with circularised *bla*_KPC_-carrying plasmids**

To infer the plasmid type that short-read contigs carrying *bla*_KPC_ genes (*n*=312) were most likely derived from, we used NUCmer (Kurtz et al. 2004) to compare them to each of 24 circularised *bla*_KPC_-carrying plasmids from the hybrid assemblies. NUCmer allows for structural rearrangements such as insertion of Tn*4401* at different locations in the same plasmid. Upon testing of this approach, we found that 98.3-100% of the sequence in each short-read contig could be aligned to the complete *bla*_KPC_-carrying plasmid from the same isolate (although in many cases to other *bla*_KPC_-carrying plasmids too). For all other comparisons, we thus required ≥98% of the short-read contig sequence to be aligned to a complete *bla*_KPC_-carrying plasmid for it to be deemed a match.

**Sequence comparisons of ColRNAI and backbone II plasmids with and without *bla*_KPC_ genes**

To infer whether *bla*_KPC_-carrying plasmids are being imported from outside of the ST258/512 lineage, or whether they result from mobilisation of *bla*_KPC_ onto existing plasmids, we compared plasmids of the same type with and without *bla*_KPC_ genes. First, we compared two ColRNAI plasmids, with and without *bla*_KPC_, from EuSCAPE_AT027 and EuSCAPE_UK048, respectively. We found high structural homology and 0 SNPs across the entire aligned sequence (**Supplementary Figure 10**). Second, we compared two backbone II plasmids, with and without *bla*_KPC_, from EuSCAPE_IT222 and EuSCAPE_IT266, respectively. We found three SNPs over 195.9kb of aligned sequence, as well as three insertions (including of Tn*4401*) in the first plasmid relative to the second (**Supplementary Figure 11**). The overall high conservation between these plasmid pairs supports movement of *bla*_KPC_ onto pre-existing plasmids in these cases.

**Additional authors**

The following are additional authors from the ESCMID Study Group for Epidemiological Markers (ESGEM):

Guido Werner^1^, Frederic Laurent^2^, Joao Carrico^3^, Edward J. Feil^4^, Ana Budimir^5^, Hajo Grundmann^6^

^1^Department of Infectious Diseases, Robert Koch Institut Wernigerode, Wernigerode, Germany, ^2^Institut for Infectious Agents, CHU de Lyon - Hôpital de la Croix-Rousse, Lyon, France, ^3^Molecular Microbiology and Infection Unit, Universidade de Lisboa, Lisbon, Portugal, ^4^Milner Centre for Evolution, University of Bath, Bath, UK, ^5^Department of Clinical and Molecular Microbiology, University Hospital Centre Zagreb, Zagreb, Croatia, ^6^Institute for Infection Prevention and Hospital Epidemiology, Medical Center – University of Freiburg, Freiburg, Germany
